# B Lymphocyte Protein Factories produced by Hematopoietic Stem Cell Gene Editing

**DOI:** 10.64898/2026.01.16.699998

**Authors:** Harald Hartweger, Chiara Ruprecht, Kai-Hui Yao, Philippe Laffont, Gabriella Lima Dos Reis, Pengcheng Zhou, Thomas Hägglöf, Laurine Binet, Maximilian Loewe, Jun P. Hong, Tianli Xiao, Esen Sefik, Brianna Hernandez, Anna Gazumyan, Mila Jankovic, Michael S. Seaman, Giulia Costa, Sean A. Nelson, Jordan Clark, Sachie Kanatani, Patrick C. Wilson, Florian Krammer, Elena A. Levashina, Jean-Philippe Julien, Hedda Wardemann, Photini Sinnis, Leonidas Stamatatos, Richard A. Flavell, Michel C. Nussenzweig

## Abstract

Long-term *in vivo* production of therapeutic proteins and development of vaccines that elicit protective levels of broadly neutralizing antibodies (bNAbs) against major pathogens face challenges. Here we report on an alternative gene-editing approach using small numbers of hematopoietic stem and progenitor cells (HSPCs) to direct long-term, high-level expression of antibodies or cargo proteins. Edited B lymphocyte offspring can be activated by cognate antigen to undergo clonal expansion and develop into specific antibody or cargo protein-synthesizing plasma cells. These cells produce long-lasting, therapeutic levels of serum antibody against HIV-1 or malaria and an anti-influenza virus bNAb that mediated universal protection from heterologous lethal challenge. Our data provide a paradigm for cell therapy approaches to prevent or treat disease using self-amplifying B cell protein factories.

## Main Text

Cells of the adaptive immune system display features that make them exceptionally good candidates for gene therapy approaches. These include longevity, circulation between tissues and the blood, and ability to respond to specific antigens that trigger effector functions and clonal expansion. These features account for the success of genetically engineered T cell-based therapies for cancer.

B lymphocytes share many of these features with T cells but are specialized to develop into protein producing long-lived plasma cells. Individual plasma cells can produce and secrete up to 10^4^ molecules of antibody per second (*1–4*) and they can survive in the bone marrow for years accounting for lifelong protection by some vaccines. These properties make B lymphocytes attractive targets for protein replacement therapies and long-term production of therapeutic or protective antibodies including antibodies that are difficult to elicit by vaccination.

Several investigators have shown that they can engineer mature B cells to produce specific antibodies and proteins (*5–22*). However, immunization based-approaches using these cells were relatively short lived and unable to consistently maintain specific protein or antibody production for long periods of time making them less desirable candidates for therapeutic antibody or protein delivery (*9, 10, 13, 15, 16, 20, 22*). Alternatively, transfer of in vitro differentiated plasma cells from edited mature B cells has shown good grafting and longer antibody expression but titers still wane over weeks to months and cannot be boosted other than by redosing (*5, 10, 14, 23–26*). While some of these hurdles may be overcome, other studies suggested hematopoietic stem and progenitor cells (HSPCs) which give rise to all blood cells, may be a better source for edited B cells (*27–31*).

Here we investigated whether very low numbers of HSPCs genetically engineered to express specific antibodies, could develop into B cells that generate long lived immune responses upon immunization and produce high levels of boostable, protective broadly neutralizing antibodies against pathogens for the lifetime of a mouse.

## Results

### Engineering murine HSPCs for optimized antibody production

While HSPCs give rise to B cells, the antibody encoding IgH and IgK loci are not expressed before commitment to the B cell lineage. To optimize antibody expression, we therefore used primary, mature, murine B cells to test numerous constructs targeting the *IgH* and *IgK* loci starting with previous designs (Fig. S1) (*9, 10, 12, 13, 15, 16, 18, 32*). B cells were gene edited using CRISPR/Cas9 sgRNA ribonucleoproteins and constructs delivered as recombinant adeno-associated virus serotype 6 (AAV6). The best construct targeted the first intron of the *IgH* locus, 3’ of Ighj4 (Fig. 1A). This construct contains a polyadenylation site to prevent expression of any *IgH* gene that undergoes productive variable-diversity-joining (V(D)J) recombination followed by an endogenous IgV_H_ insulator derived from Ighv9-4 (*33*) to focus the endogenous *IgH* intronic enhancer on the engineered *Ighv* gene promoter (Fig. 1A). The engineered *Ighv* promoter drives expression of a complete antibody light chain gene followed by a Furin-P2A cleavage site and the corresponding *IgV_H_*. This configuration directs expression of a complete Ig light and heavy chain pair and allows for class switch recombination to all Ig isotypes in secreted and membrane bound forms. Cell culture and gene editing conditions for hematopoietic stem progenitor cells (HSPCs) and long-term hematopoietic stem cells (LT-HSCs) were established by targeting mNeonGreen (mNeon) into the *B2M* locus (Fig. S2) (*34–36*).

**Fig. 1.**
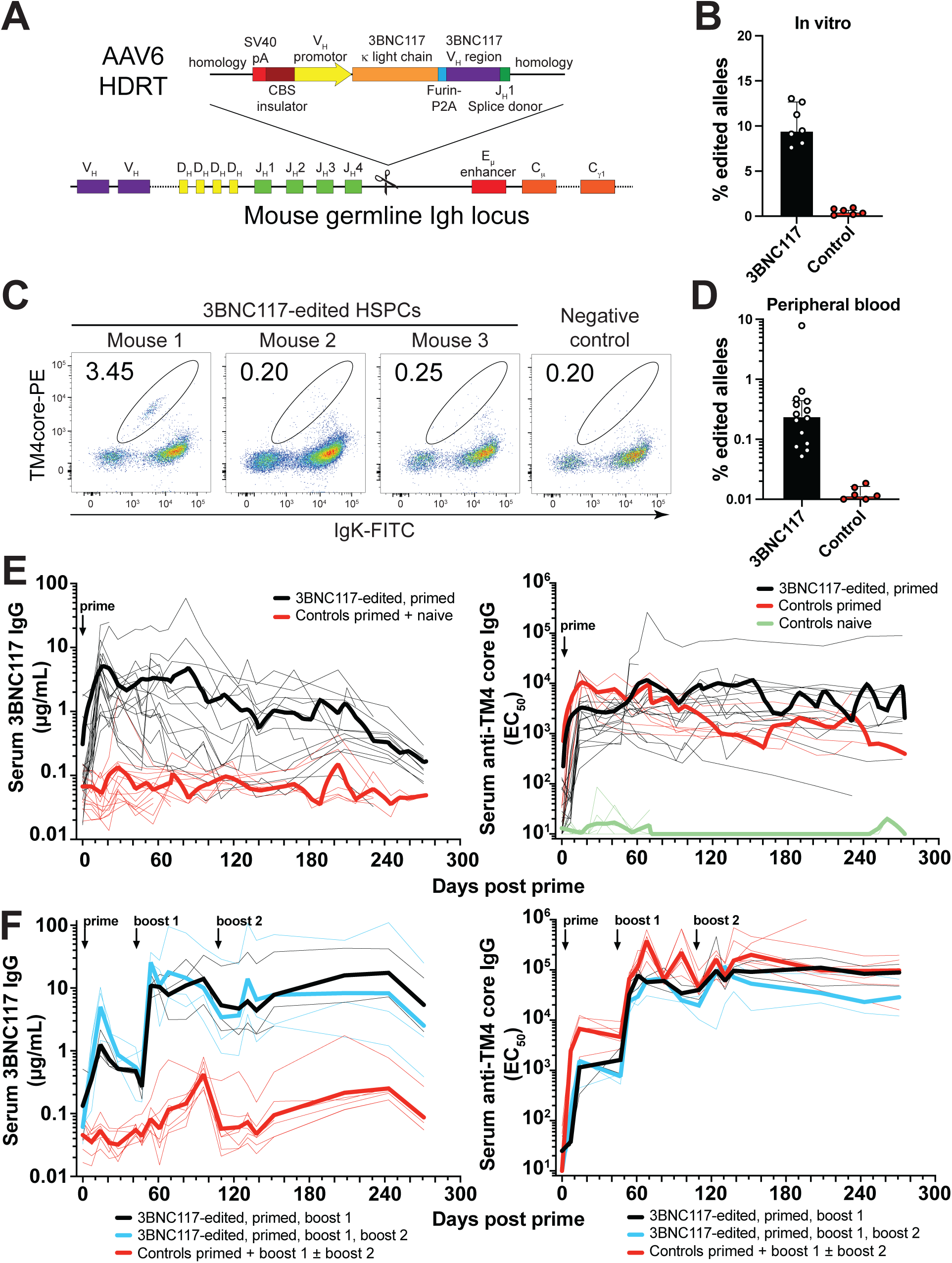
Boostable long-term expression of anti-HIV bNAb 3BNC117 through gene-edited HSPCs. **(A)** Gene editing strategy to edit antibody 3BNC117 into the mouse IgH locus after the J_H_4 intron in HSPCs. Single guide RNAs (sgRNA)/Cas9 ribonucleoproteins (RNPs) target the J_H_4 intron. The homology-directed repair template (HDRT) was packaged into AAV6. Homology arms of 661 nt (5’) or 300 nt (3’) flanked an SV40 poly A (pA) sequence, a CTCF-binding site (CBS) genetic insulator element containing the mouse Vh9-4 promoter driving the expression of a leader sequence-containing, human 3BNC117 V_L_ region fused to the mouse 𝜅 light chain constant region. A furin-cleavage site, a glycine-serine-glycine (GSG)-linker, and a P2A self-cleaving oligopeptide separate the light chain from another leader sequence, the human 3BNC117 V_H_ region which is followed by the mouse J_H_1 splice donor to allow splicing into 3’ endogenous constant regions. **(B)** Gene-editing efficiency *in vitro* in HSPCs at the time of cell transfer by digital droplet PCR measuring target locus integration using an in-out primer strategy. Controls composed of unedited and irrelevant edited samples. Pooled data from 2 independent experiments with 2–3 biological replicates and 1–2 technical replicates is shown, each dot representing a technical replicate. Bars indicate median ± interquartile range. Representative of 7 independent experiments with different HDRTs targeting the same site. **(C)** Gene-editing efficiency by flow cytometric cognate antigen binding (TM4 core) in peripheral blood donor B cells (CD45.2^+^ B220^+^) of mice 6–7 weeks after transplantation of congenic CD45.2^+^ 3BNC117-edited mouse HSPCs into CD45.1^+^ recipients. 3 representative mice from 2 independent experiments are shown. Representative of 7 independent experiments. Numbers indicate percentage of cells within gate of all cells plotted. Upstream gating as in Fig. S3A using different fluorophores. (**D**) Gene editing efficiency by ddPCR among CD45.2^+^ donor B cells or control CD45.1^+^ recipient B cells sorted from peripheral blood 6–8 weeks after transplantation of congenic 3BNC117-edited HSPCs. Gating see Fig. S3A. Data pooled from 3 independent experiments, each dot representing one mouse. Bars indicate median ± interquartile range. (**E**) 6–38 weeks after transplantation of congenic 3BNC117-edited HSPCs, mice were immunized with TM4 core, a cognate antigen for 3BNC117 and serum samples taken longitudinally, life-long or up to 273 days after immunization. Serum antibody responses were measured by anti-idiotypic 3BNC117 IgG ELISA (left panel) and total anti-TM4 core IgG ELISA (right panel). Data combined from 6 independent experiments. See Data S1 for details and Fig. S4A late primed mouse mentioned in the main text and highlighted in the same data. Thin lines indicate individual mice, thick lines indicate a group’s LOWESS local regression. (**F**) As in (E) but 7–8 weeks after transplantation indicated groups of mice were boosted once or twice with TM4 core after 47 and 110 days. Pooled data from 2 independent experiments. See Fig. S4B for low chimerism mouse mentioned in the main text and highlighted in the same data. Thin lines indicate individual mice, thick lines indicate a group’s geometric mean.

### Antibody gene-edited murine HSPCs produce long lasting, boostable immune responses

To determine whether engineered stem cells would produce B cells, we inserted 3BNC117 a broadly neutralizing anti-HIV-1 monoclonal antibody into the *IgH* locus of mouse CD45.2^+^ HSPCs (Fig. 1A, B) (*37*). 6–7 weeks after transplantation into irradiated, congenic CD45.1^+^ wildtype mice, flow cytometry and digital droplet PCR (ddPCR) assays of peripheral blood showed variable numbers of 3BNC117 expressing B cells ranging from undetectable to 4 % (Fig. 1C, D, Fig. S3A). In comparison, T cells stemming from the same HSPCs showed higher rates of editing than B cells suggesting that editing rendered some IgH loci non-functional which would interfere with B but not T cell development (Fig. S3B-E). Despite the low number of edited cells, all mice immunized with the cognate HIV-1 antigen TM4 core, produced high levels of 3BNC117 IgG as measured by anti-idiotype enzyme-linked immunosorbent assay (ELISA) whilst also producing a concurrent, polyclonal antibody response (geometric mean 3.48 µg/mL 3BNC117, range 0.48–22.32 µg/mL, Fig. 1E, Data S1).

Following immunization, 3BNC117 levels declined slowly over the lifespan of the mice for up to 273 days including mice that were immunized 38 weeks after transplantation (Fig. 1E, left and Fig. S4A). When boosted with a second dose of HIV-1 antigen 6 weeks after the prime, 3BNC117 levels increased by an order of magnitude and were maintained for at least 9 months (geometric mean 16.72 µg/mL, Fig. 1F). A further boost neither increased anti-HIV-1 IgG nor 3BNC117 IgG levels indicating saturation of the antibody response (Fig. 1F). Similar results were also obtained in mice that were reconstituted after sublethal irradiation and showed low levels of chimerism or mice boosted 10 weeks after the prime (Fig. S4B and C). The results of the anti-idiotype ELISA were generally confirmed by pseudotype virus neutralization assays against 4 different clades of HIV-1. However, the antibodies were slightly less effective at neutralizing than predicted by the anti-idiotype ELISA, likely due to low level anti-human idiotype antibody cross-reactivity with polyclonal mouse IgG (Fig. S4D, Data S2). Low levels of both polyclonal anti-TM4core IgM and IgA as well as 3BNC117 IgM and IgA were also present after immunization (Fig. S5). The levels of antibody in circulation after booster immunization were equivalent to levels that protect against infection in primates and similar to neutralizing serum Ig levels observed in the individual 3BNC117 donor (*37, 38*).

### Small numbers of edited HSPCs suffice to induce antibody responses

The relatively high-level serologic responses in mice with suboptimal chimerism suggested that small numbers of edited HSPCs were sufficient to produce antibody responses (Fig. S4B). To determine the number of edited HSPCs required to produce B cells that support humoral immune responses, we performed competitive, mixed chimera experiments in which the number of transferred gene-edited HSPCs was titrated against a constant number of wildtype bone marrow cells (Fig. 2A-C). The fraction of edited B cells in peripheral blood was determined after reconstitution by ddPCR 1–2 days before immunization. At the high dose of cultured HSPCs (2.7 × 10^5^) 0.15 % of the alleles in circulating B cells carried the edited antibody (Fig. S6A-C). The number of edited B cells in circulation dropped to undetectable in mice receiving fewer than 10^4^ cultured HSPCs (Fig. S6C). Initial 3BNC117 responses to the prime with the cognate HIV-1 antigen TM4 core were directly proportional to the number of transferred HSPCs, with little or no measurable response at doses below 10^4^ cells (Fig. 2A and B, Fig. S6D, E). However, mice receiving as few as 370 cultured HSPCs, of which on average only 29 cells were edited, produced a geometric mean of 0.11 µg/mL of 3BNC117 IgG 7 days after the boost 1. (Fig. 2B, Fig. S6D, E). A second boost replenished antibody levels but did not increase them over peak levels after the first boost (Fig. 2B). After boosting, 3BNC117 IgG levels were similar in groups receiving 29-262 edited cells/mouse suggesting these responses represent the minimum measurable response. We conclude that even very small numbers of edited HSPCs are sufficient to support development of B cells capable of producing antibody responses to antigenic challenge and these levels can be maintained by booster immunizations.

**Fig. 2.**
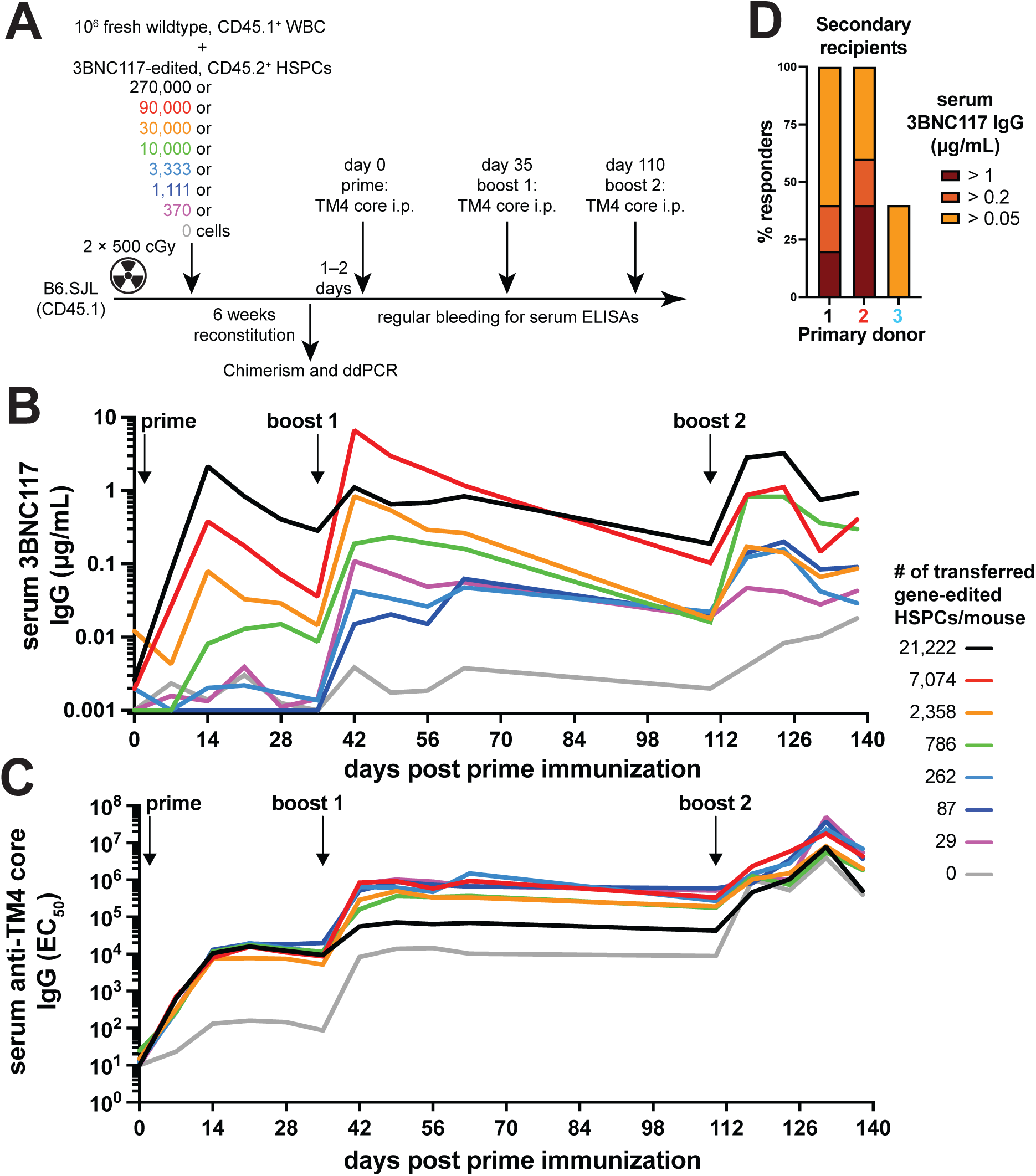
Small numbers of edited HSPCs produce edited B cells and antibodies. (**A**) Schematic experimental set up of competitive mixed bone marrow chimeras. B6.SJL mice were lethally irradiated and transplanted with a mix of 10^6^ fresh, wildtype CD45.1^+^ whole bone marrow cells (WBC) and a 3-fold serial dilution series of 3BNC117-edited CD45.2^+^ HSPCs. 6 weeks after transplantation mice were analyzed for chimerism and editing was determined by ddPCR of sorted peripheral blood CD45.2^+^ B cells. Mice were then primed and boosted after 5 weeks with 3BNC117-cognate antigen TM4 core and bled longitudinally for ELISAs. (**B**) 3BNC117 serum levels by anti-3BNC117 idiotype IgG ELISA and (**C**) serum anti-TM4 core IgG in the mixed bone marrow chimeras after TM4 core immunization. Geometric mean of n = 2-4 mice/group is plotted. See Fig. S6A-D for details. (**D**) 18-19 weeks after primary transplantation of congenic 3BNC117-edited HSPCs, secondary transplantation of CD19-depleted bone marrow was performed. 11 weeks after secondary transplantation, secondary recipients were immunized and boosted after another 5 weeks with TM4 core. Bars show the percentage of secondary recipient mice responding to TM4 core immunization by production of 3BNC117 IgG at any time point above the indicated threshold. 5 secondary recipients per donor were analyzed. 3 different primary recipients were used as donors (Primary donors) for secondary recipients. Primary recipients from 2 independent experiments were used. See Fig. S6F-K for details.

### Long-term hematopoietic stem cells can be a source of edited B cells

The transplanted HSPC cultures contained a mixture of long-term (LT-)HSPCs and more differentiated progenitors (Fig. S2E). To determine whether *bona fide* LT-HSCs could be the source of edited B cells, we performed secondary transplantation experiments using B cell depleted bone marrow cells from primary recipients 18–19 weeks after primary transplant (Fig. 2D, Fig. S6F-K). Fifteen secondary recipients received B cell-depleted bone marrow from 3 donors and were immunized and boosted with the HIV-1 cognate antigen 11 weeks after the secondary transplantation. The resulting 3BNC117 antibody responses were considerably weaker than in primary cohorts, probably due to the dilution of edited cells, nevertheless, of 15 mice 3 showed no response, 7 showed low but detectable responses, 3 responded by producing 0.2–1 µg/mL 3BNC117 IgG, 1 produced 4 µg/mL and another 48 µg/mL after the boost (Fig. 2D, Fig. S6K). In summary, a few hundred edited, transferred HSPCs could give rise to functional B cells that can be derived from LT-HSCs indicating that edited B cells in these mice would be produced for the lifetime of the organism.

### Cellular dynamics and cargo protein expression

To facilitate examination of cellular differentiation to germinal center (GC) B cells, memory B cells and plasma cells, we introduced a fluorescent cargo protein, mNeonGreen (mNeon), along with 3BNC117 into the *IgH* locus (Fig. 3A). Edited HSPCs were transplanted into congenically marked mice and immunized with the cognate HIV-1 antigen TM4 core (Fig. 3B, and Fig. S7A). The fraction of circulating B cells co-expressing both mNeon and 3BNC117 at baseline was similar to engineered B cells expressing 3BNC117 alone (Fig. S7A, B). Moreover, immunization produced 3BNC117 serum levels that matched experiments with the original construct (Fig. S7C). mNeon expressing B cells entered and underwent clonal expansion in GCs as attested to by nearly 2 log enrichment in draining lymph node GCs compared to the naïve compartment, 15.4 % vs. 0.18 % respectively peaking at day 14 after which responses subsided with typical kinetics (Fig. 3C, D and Fig. S7D, E). Accordingly, mNeon-3BNC117 expressing cells were also found in splenic and bone marrow plasma cell and switched memory B cell compartments (Fig. 3E-H and Fig. S7F-M). In summary, edited B cells arising from HSPCs were appropriately responsive to immunization and capable of producing both the edited antibody and a cargo protein.

**Fig. 3.**
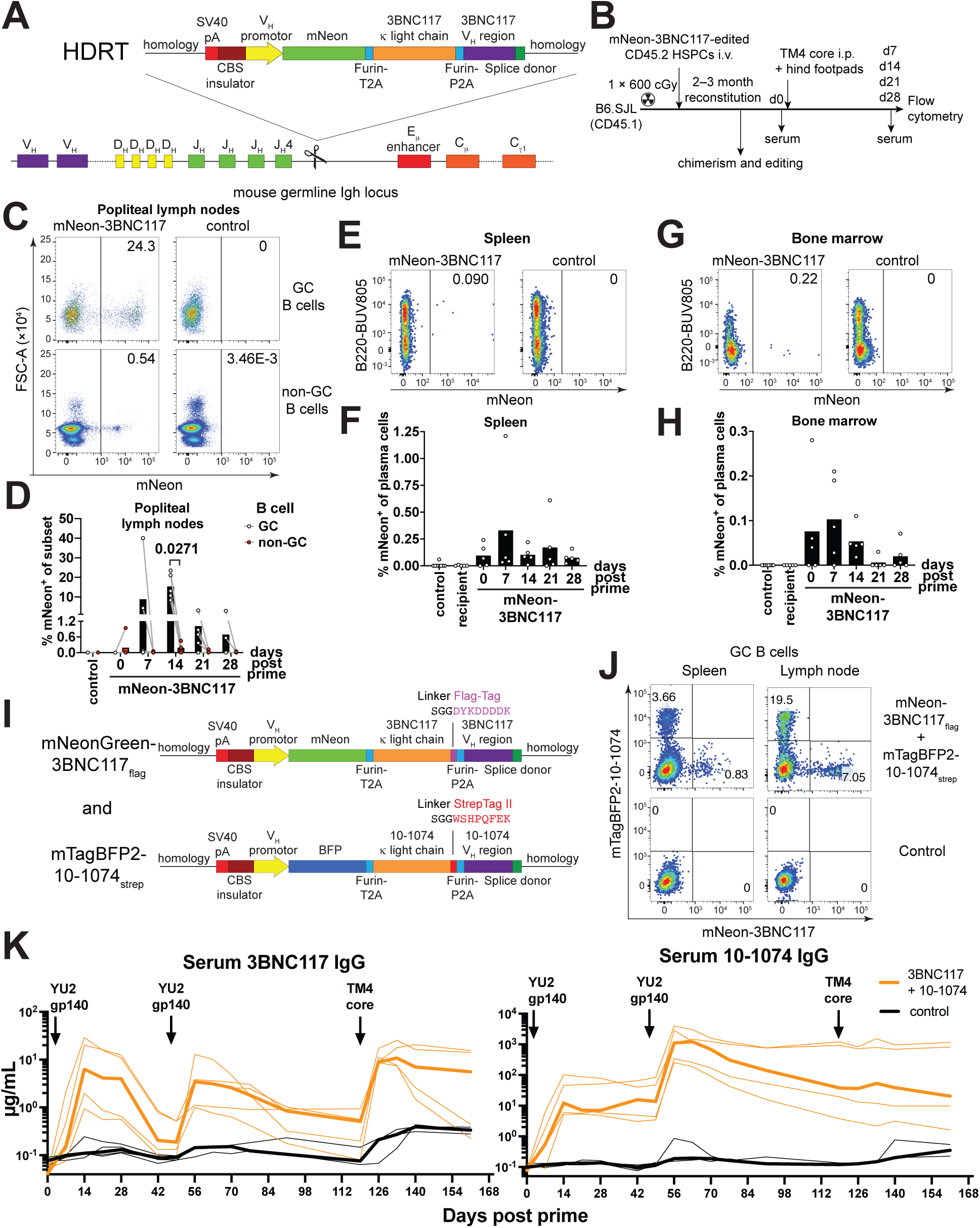
Dual antibody and cargo protein expression by edited HSPCs. **(A)** Gene editing strategy to edit mNeonGreen (mNeon) and 3BNC117 into the mouse *IgH* locus at the J_H_4 intron in HSPCs. Editing as in Fig. 1A with the following modifications: Homology arms of 558 nt (5’) or 270 nt (3’), mouse Vh9-4 promoter driving the expression of mNeon separated by a furin-cleavage site, a glycine-serine-glycine (GSG)-linker and a T2A self-cleaving oligopeptide from the downstream antibody 3BNC117. **(B)** Layout of the experiment shown in (C-H). B6.SJL were sublethally irradiated and i.v. injected with mouse HSPCs gene edited as in (A). After transplantation mice were analyzed for chimerism and gene editing and then immunized with cognate antigen TM4 core. Serum was sampled longitudinally and the mice sacrificed for analysis by flow cytometry 14 days after immunization. (**C**) Flow cytometric stain of popliteal lymph nodes 14 days after immunization gated on CD45.2^+^ GC or CD45.2^+^ non-GC B cells (gating see Fig. S7D). (**D**) Quantification of (C) showing enrichment of mNeon-3BNC117 B cells in germinal centers over 28 days after cognate antigen immunization among total cells of a subset. Each dot represents a mouse; bars indicate mean, connecting lines indicate paired data from the same mouse. Pool of 4 independent experiments with 2 independent experiments per time point. (**E**) Flow cytometric stain of spleen 14 days after immunization gated on CD45.2^+^ plasma cells (gating see Fig. S7F). (**F**) Quantification of (E) showing differentiation of mNeon-3BNC117 B cells into splenic plasma cells over 28 days after cognate antigen immunization among total plasma cells. Each dot represents a mouse; bars indicate mean. Pool of 4 independent experiments with 2 independent experiments per time point. (**G**) Flow cytometric stain of bone marrow 14 days after immunization gated on CD45.2^+^ plasma cells (gating see Fig.S7H). (**H**) Quantification of (G) showing differentiation of mNeon-3BNC117 B cells into bone marrow plasma cells over 28 after cognate antigen immunization among total plasma cells. Each dot represents a mouse; bars indicate mean. Pool of 4 independent experiments with 2 independent experiments per time point. (**I**) Gene editing strategy as in (A) but adding a Flag-tag or StrepTag II to the C-terminus of the light chain and tagging antibody 10-1074 with mTagBFP2 fluorescent protein. Cells were edited separately and mixed for transplantation. (**J**) Flow cytometric analysis of edited B cell recruitment into germinal centers 14 days after immunization of mice receiving HSPCs edited as in (I), reconstituted for 6 weeks and immunized with TM4core and YU2-gp140 i.p. and into footpads. Gated on GC B cells. Representative of 3 mice. (**K**) 3BNC117 and 10-1074 IgG serum levels by anti-idiotype ELISAs in mice receiving a mix of separately edited untagged 3BNC117 and 10-1074 edited HSPCs or unedited C57BL/6J control animals after reconstitution and YU2-gp140 prime and boost and TM4 core secondary booster immunization. Representative of 2 independent experiments. Thin lines indicate individual mice, thick lines indicate a group’s median. Paired, one-tailed T-test P value in (D) is shown. Numbers in flow cytometric plots indicate percentage of cells within gate of all cells plotted.

To characterize development of edited B cells, we created chimeras using allotype heterozygous IgH^a^/IgH^b^ CD45.2 donor HSPCs. The cells were edited with an mNeon-3BNC117 construct and reconstituted IgH^b^ CD45.1 recipient mice (Fig. S8A). Edited pre-B cells were detected in the bone marrow and accumulated in the mature B cell compartment with no discernible blocks during B cell development (Fig. S8B, C). IgH allotype analysis suggested intact allelic exclusion of the heavy chain and consequently mostly monoallelic gene-editing (Fig. S8D-G). We detected a strong bias (up to 94 %) in either IgH^a^ or IgH^b^ allele usage in edited B cells between mice compared to their polyclonal repertoire (∼50 %) (Fig. S8H, I) suggesting an oligoclonal source with few edited HSCs giving rise to edited B cells.

### Combination of antibodies

To determine whether combinations of antibodies could be produced simultaneously, we engineered two sets of HSPCs individually, one with anti-HIV-1 antibody 3BNC117 and one with anti-HIV-1 antibody 10-1074 (*39*). These antibodies target two different non-overlapping epitopes on the HIV-1 viral envelope protein. Mice were reconstituted with a mix of both sets of separately gene-edited HSPCs and immunized with a single cognate high affinity HIV-1 antigen for both antibodies (YU2 gp140). 3BNC117 or 10-1074 expressing B cells were both recruited into germinal centers and produced high levels of the two antibodies (Fig. 3I-K, Fig. S9A, B). Serum 10-1074 was expressed at levels up to a median of 1.2 mg/mL two weeks after boost. Additionally, 3BNC117 antibody levels were specifically increased in a second boost using an antigen (TM4 core) with high affinity for 3BNC117 but not 10-1074 (Fig. 3K, Fig. S9C, D). In summary, HSPCs engineered to express different antibodies could be used in combination *in vivo*.

### Human HSPC antibody gene editing

To determine whether human HSPCs could also give rise to B cells, we adapted the mouse constructs for use with human cells (Fig. S10) and optimized the conditions by targeting mNeon into the human B2M locus (Fig. S11) (*9, 18, 40–43*). Human HSPC editing was far more efficient than for mouse HSPCs. We achieved up to 56–75 % targeting efficiency for the B2M locus in 4 different donors of human fetal liver-derived CD34^+^ HSPCs (Fig. 4A, Fig. S11B-G). The *IgH* locus of the human HSPCs was edited with anti-HIV-1 antibodies 3BNC117 or 10-1074 achieving 40 % editing efficiency (Fig. 4B). Edited human HSPCs carrying 3BNC117 or B2M tagged with mNeon were transplanted into immunodeficient MISTRG6 mice (*44*) (Fig. 4B-F, Fig. S12). 3BNC117 expressing B cells represented on average 21 % of the human CD19^+^ cells in circulation of the reconstituted mice (Fig. 4E, F). Improved engraftment in this model may be artificially high due to host immunodeficiency. Nevertheless, the data show that human HSPCs can be edited efficiently and retain the potential to differentiate into B cells *in vivo*.

**Fig. 4.**
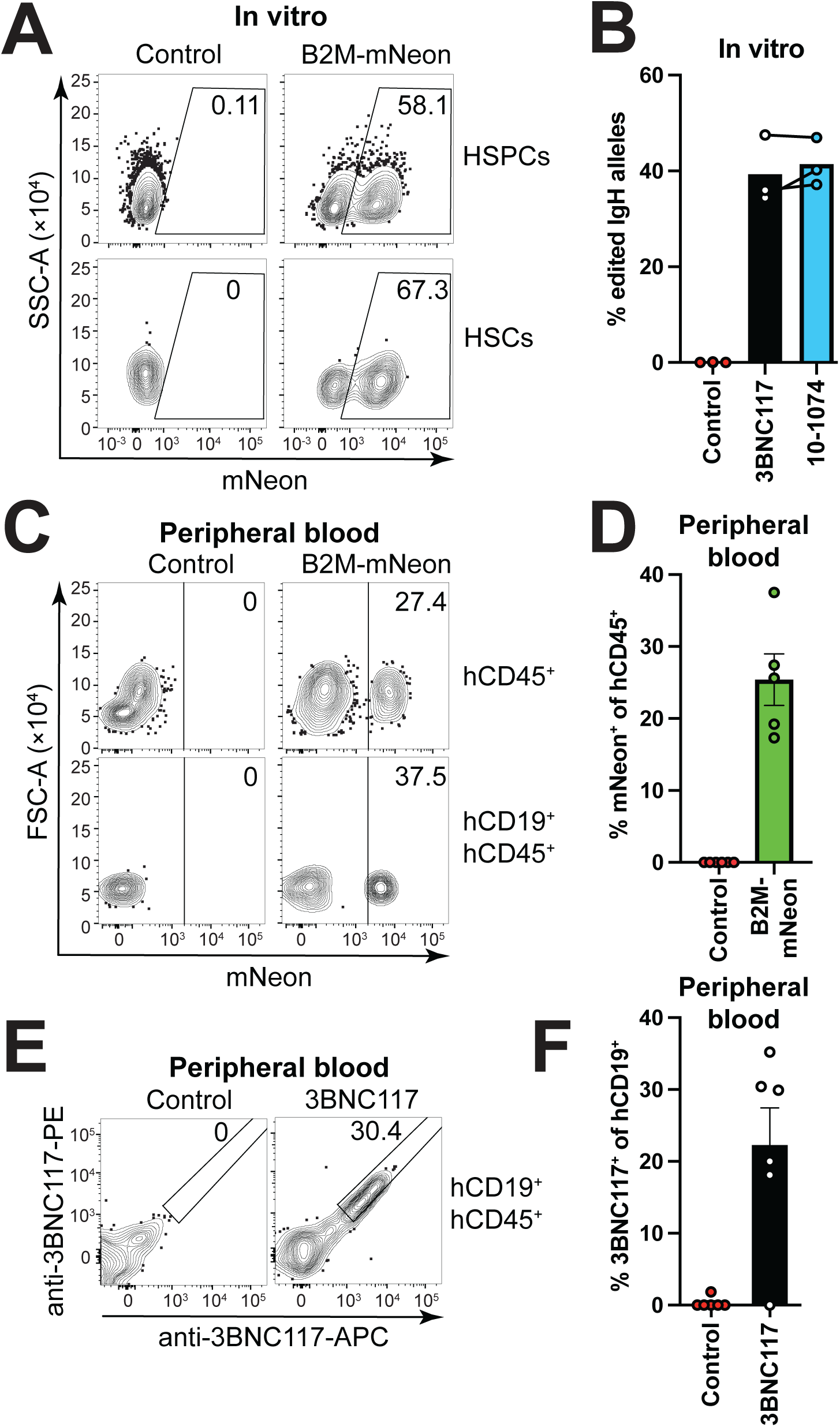
**Anti-HIV-1 antibody expressing human B cells**. (**A**) Human fetal liver CD34^+^cells were cultured for 2 days, then gene-edited to express mNeon from the B2M-locus using Cas9 RNP and an HDRT packaged into recombinant AAV6. Flow cytometric analysis of mNeon expression in primitive HSPCs (HSPCs, CD34^+^, CD38^-^) and LT-HSC phenotype cells (HSCs, CD34^+^, CD38^-^, CD90^+^, CD45RA^-^, CD201^+^) 18 h after transduction is shown. Gated as in Fig. S11C. (**B**) Cells were edited as in (A) but inserting antibodies 3BNC117 or 10-1074 into the *IgH* locus using CBS-containing promoter-driven constructs (see Fig. S10 for details). ddPCR was performed on gDNA of cultured cells detecting edited *IgH* alleles using an in-out primer strategy and normalized to a genomic reference PCR. Samples were cultured for 24 h then edited and harvested after another 18 h. Controls are irrelevantly edited cells. Bars indicate mean. Each dots represents a technical replicate from a different donor. Identical donors for 3BNC117 and 10-1074 edited cells are connected by a line. Representative of 3 independent experiments. (**C**) Flow cytometric analysis of mNeon expression among total human hematopoietic cells (hCD45^+^) or human B cells (hCD19^+^, hCD45^+^) of peripheral blood of MISTRG6 mice 6 weeks after transplantation of gene-edited human CD34^+^ fetal liver cells. Controls mice received irrelevantly edited cells. Gated as in Fig. S12. (**D**) Quantification of (C). Each dots represents a mouse, bars indicate mean ± SEM. Representative of 3 independent experiments. (**E**) Flow cytometric analysis of 3BNC117 expression using an anti-3BNC117-idiotype stain among human B cells (hCD19^+^, hCD45^+^) of peripheral blood of MISTRG6 mice 8 weeks after transplantation of gene-edited human CD34^+^ fetal liver cells. Control mice received irrelevantly edited cells. Gated as in Fig. S12. (**F**) Quantification of (E). Each dot represents data from an individual mouse, bars indicate mean ± SEM. Representative of 3 independent experiments. Numbers in flow cytometric plots indicate percentage of cells within gate of all cells plotted. Upstream gating for (B) and (D) as in Fig. S3A using different fluorophores.

### Induction of high titer anti-Malaria parasite antibodies

To determine whether other antibodies could also be produced by edited, murine B cells we designed targeting constructs to direct expression of anti-*Plasmodium falciparum* circumsporozoite protein (PfCSP) antibodies 317 (*45*), 2541 (*46*) and iGL-CIS43.D3 (*47*) or murine control antibody B1-8^hi^ specific for the hapten (4-hydroxy-3-nitrophenyl)acetyl (NP) (*48, 49*) (Fig. 5). To enable specific detection of these antibodies in the absence of anti-idiotypic reagents, we incorporated a Flag-tag into the carboxyl terminus of their Ig light chains (*50*) (Fig. S13A-C).

**Fig. 5.**
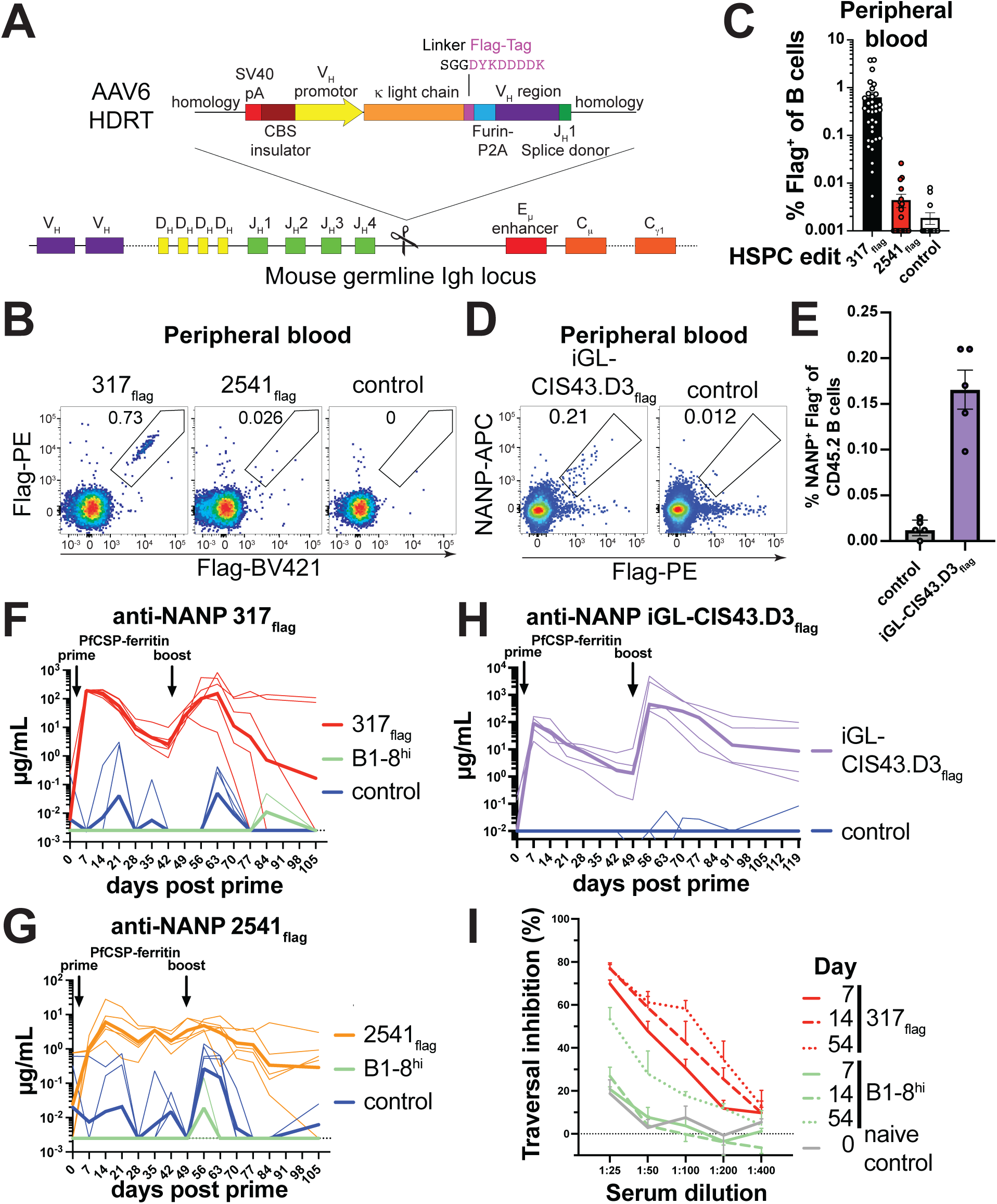
Anti-*Plasmodium falciparum* antibody expression *in vivo*. (**A**) Gene editing strategy to edit light chain flag-tagged antibodies 317, 2541 or iGL-CIS43.D3 into the mouse *IgH* locus. (**B**) Flow cytometric analysis of flag-tagged antibody surface expression on CD45.2^+^ B cells from peripheral blood of mice 5.7 weeks after receiving congenic HSPCs edited with the indicated antibody. A non-flag tagged B1-8^hi^ antibody was used as control. (**C**) Quantification of (B). Each dot represents a mouse, bars indicate mean ± SEM (**D**) Flow cytometric analysis of flag-tagged, NANP peptide-binding antibody surface expression on CD45.2^+^ B cells from peripheral blood of mice 6 weeks after receiving congenic HSPCs edited with iGL-CIS43.D3_flag_. Controls were edited with irrelevant antibodies. (**E**) Quantification of (D). Each dot represents a mouse, bars indicate mean ± SEM. Data from 1-4 experiments per antibody for (B-E). (**F**) 10 weeks after receiving 317_flag_ or control B1-8^hi^ edited-HSPCs or C57BL/6J wildtype control mice were immunized with PfCSP-ferritin nanoparticle immunogen and anti-NANP peptide flag-tagged serum antibody responses were measure by ELISA. Thick lines indicate geometric mean and thin lines indicate individual mice. (**G**) as in (F) but for 2541_flag_. (**H**) as in (F) but for iGL-CIS43.D3_flag._ (**I**) Inhibition of *Plasmodium falciparum* hepatocyte traversal by serially diluted pooled mouse sera from (F). Mean + SEM from three independent biological experiments is shown. Numbers in flow cytometric plots indicate percentage of cells within gate of all cells plotted.

In mice, transplanted 317_flag_, 2541_flag_, iGL-CIS43.D3_flag_ anti-PfCSP and control antibody B1-8^hi^edited HSPCs produced similar numbers of B cells as the anti-HIV-1 edited HSPCs as measured by ddPCR and flow cytometry (Fig. 1 and 5B-E, Fig. S13D-F). Upon cognate prime-boost immunization, mice produced > 100 µg/mL 317_flag_, 1–10 µg/mL 2541_flag_ and up to 1000 µg/mL iGL-CIS43.D3_flag_ respectively (Fig. 5F-H, Fig. S13G, H). Serum from 317_flag_ and B1-8^hi^ control mice were tested for the capacity to inhibit *Plasmodium falciparum* sporozoite traversal of hepatocytes *in vitro*. In line with 317_flag_ titers, serum from 317_flag_ edited mice inhibited sporozoite traversal as early as on day 7–14 after prime and the inhibitory capacity of the sera remained stable after boost (Fig. 5F, I). Serum from B1-8^hi^ control mice showed traversal inhibition only after boost, likely due to the polyclonal response to immunization increasing over time (Fig. 5I, Fig. S13G).

### Anti-influenza bNAbs, somatic hypermutation (SHM) and protection from heterologous lethal influenza virus infection

We tested whether edited cells expressing a broadly neutralizing anti-influenza virus antibody would produce universally protective antibodies against lethal infection and undergo somatic hypermutation (SHM). Mice were reconstituted with HSPCs carrying flag-tagged MEDI8852, a broadly neutralizing anti-influenza hemagglutinin stalk antibody (*51*) (Fig. 5A, Fig. 6A, Fig. S14A). 6–8 weeks after hematopoietic reconstitution 0.1 % of B cells in peripheral blood expressed the antibody (Fig. 6 A-C, Fig. S14B). When immunized with hemagglutinin from strain A/Hong Kong/1/1968 (H3N2) (HK68), mice produced a geometric mean of 11.5 µg/mL of MEDI8852_flag_ 14 days after prime and 141 µg/mL 7 days after boost whereas control animals produced none (Fig. 6D). All mice responded to immunization as polyclonal H3 HK68-binding IgG responses were present in both groups and slightly stronger in control animals (Fig. 6E).

**Fig. 6.**
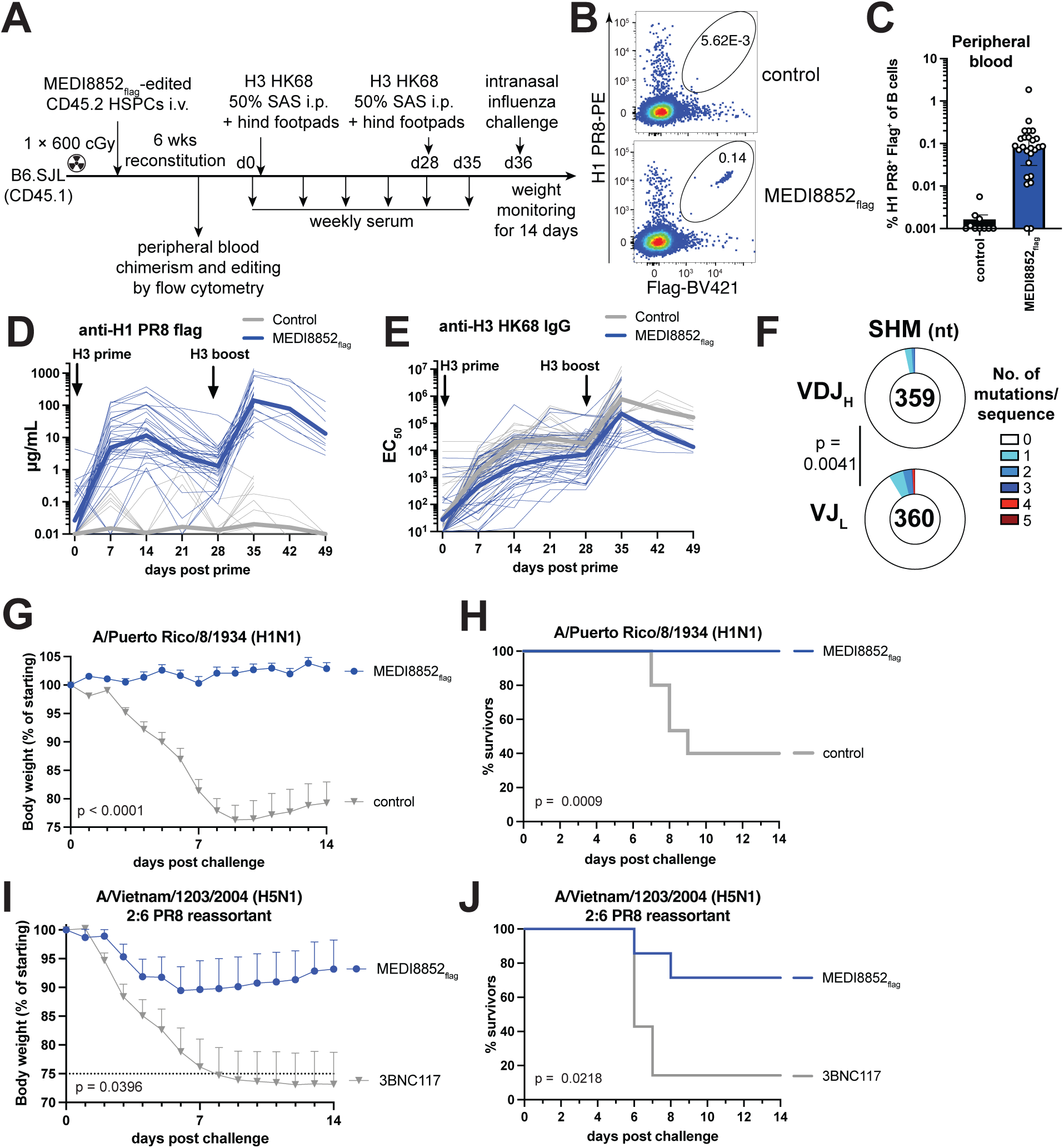
Protection from influenza infection and somatic hypermutation. (**A**) Schematic of experimental procedures in this figure. (**B**) Flow cytometric analysis of PR8 hemagglutinin (H1 PR8) binding and flag-tagged antibody surface expression on CD45.2^+^ B cells from peripheral blood of mice 5–7 weeks after receiving congenic HSPCs edited with the indicated antibody. Unedited mice were used as controls. Upstream gating as in Fig. S3A using different fluorophores. Numbers indicate percentage of cells within gate of all cells plotted. (**C**) Quantification of (B). Percentage of H1-PR8^+^Flag^+^ cells of total B cells. Each dot represents a mouse. Bars indicate median ± interquartile range. Pooled data from 4 independent experiments. Values < 0.001 set to 0.001. (**D**) ELISA quantifying flag-tagged MEDI8852 binding to H1-PR8 or (**E**) total anti-HK68 hemagglutinin (H3 HK68) mouse IgG in mouse serum of unedited non-chimeric C57BL/6J or irrelevantly edited 3BNC117 control mice, or mice receiving MEDI8852_flag_-edited HSPCs, primed 6–9 weeks after transplant with H3-HK68 and boosted after another 28 days. Thick lines indicate geometric mean and thin lines indicate individual mice. Pooled results from 6 independent experiments. Mice measured until day 35 only were used in challenge experiments in (F-I). (**F**) Number of somatic hypermutations per sequence in heavy (VDJ_H_, top) or light chain (VJ_L,_ bottom) regions of single-cell sorted Flag-tagged MEDI8852flag germinal center B cells 7 days after third immunization with H3 HK68, 322-356 days after prime. Numbers indicate total analyzed cells. Pool of 4 mice from 2 independent experiments. P-value for Fisher’s exact test is shown. (**G**) Weight loss (dots show mean + SEM) and (**H**) survival of mice upon 10×MLD_50_ (10^3^ plaque forming units, PFU) influenza PR8 virus challenge 8 days after boost (day 36). Pooled data from 2 independent experiments. n = 13–15 per group. Last weight measurement of euthanized or dead mice carried forward. Controls were C57BL/6J mice. (**I**) Weight loss (dots show mean + SEM) and (**J**) survival of mice upon challenge with 9×MLD_50_ (10^4^ PFU) A/Vietnam/1203/2004 (H5N1) PR8 reassortant with deleted polybasic cleavage site, challenged 8 days after boost (day 36). One of 2 independent experiments is shown. n = 7 per group. (F and H) P value of two-way ANOVA is shown. (G and I) P value of Log-rank test is shown.

To test if edited B cells underwent somatic hypermutation, HK68 primed mice were boosted twice and edited germinal center B cells purified from draining lymph nodes for single cell mutation analysis (Fig. 6F, Fig. S15, Data S3). Virtually all edited GC B cells bound heterologous H1 PR8 protein as determined by flow cytometry (Fig. S15A). Clonal expansion was detected as measured by the presence of identical mutated sequences (Fig. S15E, Data S3). SHM accumulated proximal to the V_H_ promoter in the VJ_L_ with lower mutation rates in the more distal VDJ_H_ (*52*) (Fig. 6F, Fig. 15B-D, F, Data S3). The overall rate of mutation was approximately 10^-4^ mutations/nt (Fig. S15D). We concluded that B cell progeny of antibody-edited HSPCs can somatically hypermutate in response to cognate immunization.

When prime and boosted MEDI8852_flag_-edited mice were challenged with a lethal dose (10^3^ plaque forming units (PFU), 10-fold mouse lethal dose 50 % (mLD_50_)) of phylogenetically distant strain A/Puerto Rico/8/1934 (H1N1) (PR8) that causes severe infection in mice, all control mice lost weight and 60 % succumbed to infection, while mice expressing MEDI8852_flag_ gained weight and all survived (Fig. 6G,H Fig. S16A,B). A separate group of MEDI8852_flag_ reconstituted mice that were immunized with H3 were challenged with an even more aggressive lethal dose (10^4^ PFU, 9 × mLD_50_) of phylogenetically distant strain A/Vietnam/1203/2004 (H5N1, 2:6 PR8 reassortant). The 3BNC117-edited control mice showed rapid weight loss with 85 % succumbing to infection, while the group expressing MEDI8852_flag_ showed only moderate weight loss, on average 10 % (p = 0.0396), and 70 % survived (p = 0.0218) (Fig. 6I, J, Fig. S16C, D). MEDI8852_flag_ titers the day before challenge correlated well with weight loss after challenge (Fig. S16E). We concluded that gene-edited HSPCs give rise to B cells that produce long lasting universally protective anti-influenza antibody responses.

## Discussion

Cell division in response to antigen is the basis of the clonal selection theory (*53*). Under physiologic circumstances rare antigen specific B cells respond to antigenic challenge by undergoing clonal expansion in GCs and producing long-lived memory and plasma cells. As few as one naïve, high-affinity, antigen-specific B cell can produce expanded clones of antibody-secreting plasma cells (*54–56*). Secondary challenge or boosting rapidly amplifies the antibody response. Our experiments show that the immune system can be co-opted to direct expression of specific antibodies and cargo proteins introduced into small numbers of HSPCs by gene editing.

Cell types such as muscle or liver have been the focus of gene therapy efforts to produce therapeutic proteins because of their abundance and relatively uncomplicated methods to induce expression, however these cells do not offer the same advantages as HSPC-derived B cells. Whereas other cell types generally produce therapeutic proteins constitutively from large numbers of cells, B cells allow induction of high therapeutic expression levels by specific antigenic challenge from very small numbers of cells. Moreover, decreased expression of a therapeutic protein by a liver or muscle cell would require additional genome engineering. In contrast, a booster immunization with an available vaccine would increase cargo protein expression in antigen-specific B cells that carry cognate antibody when the protein in question reaches subtherapeutic levels. Furthermore, for antibody therapies or disease prevention, B cells and their plasma cell progeny are uniquely positioned to produce high titers of lasting antibody responses.

Our experiments demonstrate that a single anti-influenza antibody produced through the progeny of gene-edited HSPCs achieves antibody levels that are universally protective against lethal infection with influenza A virus. However, other pathogens such as HIV-1 may require a combination of antibodies to be universally effective in prevention and therapy to suppress viral escape mutants. As demonstrated by our experiments with broadly neutralizing anti-HIV-1 antibodies, HSPCs can be edited individually with different antibodies to produce a combination that increases pathogen variant coverage suggesting a potential application as a functional cure for HIV-1 (*57–61*). Additionally, though unlikely as SHM was low, a fraction of the B cells may adapt to the mutating virus in a chronic infection setting while a constant source of unmutated bNAb would guarantee known breadth and potency.

When testing human antibodies in animal models poly- and autoreactivity to species-specific antigens may restrict development of edited B cells or reduce antibody serum half-live. Consistent with the idea that the paratope influences antibody stability, we observed differences in expression levels of various antibodies targeting the same antigen (Fig. 3K, 5F-H). Thus, antibodies selected for further translation will need careful screening including high affinity interaction with antigen (see supplementary text for a detailed discussion, and Fig 3K, S9D). Despite these caveats, the HSPC editing approach displayed high flexibility being able to produce six therapeutic antibodies against three major pathogens, two different cargos and the possibility to combine antibodies while showing low levels of SHM. These features should allow development of many different therapeutic strategies including antibodies or other proteins. A similar approach may also enable long-lived T cell responses against cancer.

Finally, the requirement for only small numbers of HSPCs to produce B cell protein factories makes this approach a prime candidate for direct *in vivo* gene-editing. Thus, even very inefficient *in vivo* approaches to editing that result in modification of only a very minor population of HSPCs would still be expected to yield an amplifiable group of B cells that produce large amounts of antibody for prevention, or therapy or cargo proteins for enzyme or protein replacement in genetically deficient individuals. Subject to such advances, this could be envisioned as a single shot approach to produce B cell protein factories *in vivo*.

## Materials and Methods

### Cloning and recombinant adeno-associated virus (AAV) production

Plasmids were synthesized by Twist Biosciences. Alternatively, genes were synthesized as gBlocks (IDT) and cloned into restriction-digested vectors by Gibson assembly using HiFi DNA assembly mastermix (NEB). Recombinant AAV6 was produced as previously described by us and a detailed protocol is available (*18*). In brief, pAAV plasmid encoding the desired antibody construct was transfected into 293AAV cells (Cell BioLabs) and simultaneously transduced with a Tetracycline enabled self-silencing adenovirus-RepCap6 (TESSA-RepCap6, Oxgene) providing all packaging genes (*62*). Supernatant was harvested and recombinant AAV precipitated using polyethylene glycol/NaCl followed by chloroform extraction and centrifugal filter unit buffer exchange and concentration. Recombinant AAV was stored at -80°C and titrated by ddPCR measuring an ITR amplicon as previously described (*18*).

### Mice

C57BL/6J, B6.SJL-*Ptprc^a^ Pepc^b^*/BoyJ and B6.Cg-Gpi1a Thy1a Igha/J were obtained from the Jackson Laboratory. B6.Cg-Gpi1a Thy1a Igha/J was bred to C57BL/6J to create IgH^a/b^ donor mice. The B1-8^hi^ (*63*) strain was generated and maintained at Rockefeller University on a C57BL/6J background. MISTRG6 mice (*44*) were generated at Yale University. Mice were housed at a temperature of 22 °C and humidity of 30–70 % in a 12 h light/dark cycle with ad libitum access to food and water. Sample sizes were not calculated a priori. Given the nature of the comparisons, mice were not randomized into each experimental group and investigators were not blinded to group allocation. Instead, experimental groups were age- and sex-matched. All experiments used 6–16 week old mice at the start of the experiment. Both male and female mice were used for experiments. All animal experiments, except MISTRG6 transplantation experiments, were performed at Rockefeller University with authorization from the Institutional Review Board and the Rockefeller University Institutional Animal Care and Use Committee. MISTRG6 transplantation experiments were performed at Yale University in compliance with Yale Institutional Animal Care and Use Committee protocols.

### Mouse B cell gene editing

Mouse B cells were gene edited as previously described (*9*) with the following modifications. CD43-depleted, Ter119-depleted splenic mouse B cells were cultured for 24 h in B cell culture medium composed of Roswell Park Memorial Institute 1640 medium (Gibco) supplemented with 10 % heat-inactivated fetal bovine serum (Hyclone), 2 mM glutamine, 1 mM sodium pyruvate, 10 mM N-2-hydroxyethylpiperazine-N-2-ethane sulfonic acid (HEPES), 55 µM β-mercaptoethanol, 1× non-essential amino acids and 1× insulin-transferin-selenium and 1× antibiotic-antimycotic (all Gibco). Cells were activated with either 50 µg/mL Lipopolysaccharide from *E. coli* O111:B4 (Sigma-Aldrich) or alternatively a cocktail of 100 ng/mL mouse MegaCD40L (Enzo), 5 µg/mL CpG ODN 2395 (Invivogen), 40 ng/mL human B cell activating factor (BAFF), 50 ng/mL mouse interleukin (IL)-2 and 50 ng/mL mouse IL-10 (all PeproTech). Per 10 uL transfection ribonucleoproteins (RNPs) were formed using 0.75 µL of 100 µM sgRNA in duplex buffer and 0.4 µL of 61 µM Cas9 (all IDT). If more than one sgRNA was used, separate RNPs were prepared and the final volumes above were split equimolarly between different sgRNAs. sgRNAs are listed in Data S4. After RNP complex formation, 9 µL buffer R containing 5×10^5^ activated B cells were added and electroporated at 1650 V 20 ms 1 pulse using the Neon Transfection System (Thermo Fisher Scientific) and transferred to 50 µL culture medium with stimulants but without antibiotic-antimycotic in a 48-well plate with or without recombinant AAV6. Recombinant AAV6 multiplicity of infections (MOIs) ranged from 0.125–1×10^6^, not exceeding 12.5 µL (25 % of culture volume). Cells were incubated at 37°C 5 % CO_2_ for 5 h before addition of 450 µL culture medium with stimulants with antibiotic-antimycotic. Cells were analyzed 2 days after electroporation.

### Human B cell gene editing

Healthy human leukapheresis samples were collected after signed informed consent in accordance with protocol TSC-0910 approved by the Rockefeller University Institutional Review Board. Peripheral blood mononuclear cells (PBMCs) were prepared, stored in liquid nitrogen. Human B cells were gene edited as previously described (*9*) with the following modifications. PBMCs thawed in a 37 °C water bath and isolated using the EasySep human naive B cell Enrichment Kit (19254; Stemcell) according to the manufacturer’s instructions. Human B cells were cultured in B cell culture medium supplemented with 100 ng/mL human MegaCD40L (Enzo), 3.33 µg/mL CpG ODN 2395 (Invivogen), 40 ng/mL human BAFF, 50 ng/mL human IL-2 and 50 ng/mL human IL-10, 10 ng/mL human IL-15 and 20 ng/mL human IL-4 (all PeproTech). Human B cells were edited and analyzed like mouse B cells described above with the following modifications: Cells were resuspended in buffer T and electroporated using the following settings 1750 V 20 ms 1 pulse. sgRNAs are listed in Data S4.

### Mouse HSPC enrichment and culture

Mouse HSPCs were obtained from bone marrow of C57BL/6J mice. All procedures were performed in phosphate-buffered saline (PBS) and cells centrifuged at 200 g for 10 min 4 °C unless otherwise indicated. Pelvic bones, femurs and tibiae were dissected and cut open at the iliac crest or proximal to the patella and bone marrow extracted by centrifugation at 10,000 g for 15 s (*64*). Red blood cells were lysed with ammonium-chloride-potassium (ACK) lysing buffer for 5 min and cells filtered through a 70 µm cell strainer and washed with PBS by centrifugation. Cells number was adjusted to 10^8^ cells/mL and cells were incubated with 12.5 µg/mL mouse FcBlock (BD Bioscience) for 15 min at 4°C under constant gentle agitation. Then, 4 µg/mL anti-Sca1-APC (BioLegend) was added and cells were further incubated for 20 min at 4 °C under constant gentle agitation. Cells were washed once in PBS and resuspended at 10^8^ cells/mL. Anti-APC microbeads (Miltenyi Biotec) were added at 20 % of the cell suspension volume and incubated for 15 min at 4 °C under constant, gentle agitation and then washed once more. Microbead-binding cells were enriched on LS columns according to manufacturer’s protocol and eluted in 5 mL PBS. Cells were washed, counted and resuspended at 2×10^5^ cells/mL in mouse HSPC culture medium composed of StemSpanTM SFEM II medium supplemented with mouse stem cell factor (SCF) (10 ng/mL), mouse thrombopoietin (TPO) (100 ng/mL), mouse Flt3-Ligand (50 ng/mL), human IL-11 (50 ng/mL) (all Peprotech) and 1× antibiotic antimycotic and cultured in 15 cm cell cultures plates (Falcon) at 37 °C with 5 % CO_2_ for 24–48 h.

### Mouse HSPC gene editing and transplantation

All steps were performed at room temperature with room temperature reagents and centrifugation was performed at 200 g for 10 min at room temperature. After 24–48 h in culture, mouse HSPCs were harvested by centrifugation. Cells were resuspended in PBS, counted and centrifugated again. Cells were then resuspended at 1.11-1.66×10^7^ cells/mL in buffer P4 (Lonza). Per 100 uL transfection ribonucleoproteins (RNPs) were formed using 7.5 µL of 100 µM sgRNA in duplex buffer and 4 µL of 61 µM Cas9 (all IDT). Typically only 1 sgRNA targeting the IgH locus was used per transfection. Sequences are available in Data S4. 11.5 µL RNP was mixed with 90 µL of cells per 100 µL transfection. Mouse HSPCs were electroporated using Lonza 4D nucleofector and kit using setting EN-100. Two 100 µL transfections were transferred to 1 mL mouse HSPC culture medium with aforementioned cytokines but without antibiotic-antimycotic in a 10 cm dish with or without recombinant AAV6. Recombinant AAV6 was used at MOI = 16,600 unless otherwise indicated, not exceeding 25 % of culture volume. Cells were incubated at 37°C with 5 % CO_2_ for 5 h before topping up the dish to 9-11 mL with mouse HSPC culture medium with stimulants with antibiotic-antimycotic and culture overnight. The next day, typically 18 h later, cells were harvested by centrifugation, washed once in PBS and 1–3×10^5^ cells/mouse, unless otherwise indicated, in 200 µL PBS were injected intravenously into congenically marked 6–14 week old B6.SJL-*Ptprc^a^ Pepc^b^*/BoyJ recipient mice that had been either lethally (2×5 Gy) or sublethally (1×6 Gy) irradiated, as indicated, on the same day. Post treatment, mice were preventatively kept on amoxicillin feed for 6-7 weeks before switching back to standard chow. Alternatively, cells were analyzed 1-2 days after electroporation unless otherwise indicated.

### Human HSPC culture and gene editing

We used human fetal livers derived HSPCs since they are the preferred source of donor cells for the MISTRG6 mouse model (*44*). Human fetal livers were procured from Advanced Bioscience Resources (ABR), Inc (Alameda) or the Human Fetal Tissue Repository (HFTR, Bronx, NY). CD34^+^ HSPCs were isolated as previously described (*65*). In brief, human fetal livers were homogenized and incubated in digestion medium (Hank’s balanced salt solution with 0.1 % collagenase IV (Sigma-Aldrich), 40 mM HEPES, 2 mM CaCl_2_ and 2 U/mL DNAse I (Roche)) for 30 min at 37 °C. Human CD34^+^ HSPCs were isolated using a CD34^+^ cell isolation kit (Stem Cell Technologies) according to the manufacturers’ instructions and gradually frozen in fetal bovine serum (FBS) with 10 % dimethyl sulfoxide (DMSO) and stored in liquid nitrogen until usage. Cells were thawed in a 37 °C water bath and washed twice in prewarmed StemSpan SFEM II medium (Stemcell Technologies) and cultured at 10^5^ cells/mL in human HSPC culture medium composed of StemSpan SFEM II medium supplemented with human SCF (100 ng/mL), human TPO (100 ng/mL), human Flt3-L (100 ng/mL), human IL-6 (20 ng/mL) (all Peprotech), 500 nM UM729, 750 nM StemRegenin 1 (both Stemcell Technologies) and 1× antibiotic antimycotic (Gibco) in 15 cm cell culture plates (Falcon) at 37 °C with 5 % CO_2_ for 24 h. Human HSPCs were edited as described for mouse HSPCs with the following modifications: Cells were resuspend in buffer P3 (Lonza) and electroporated using setting DZ-100. Recombinant AAV6 was used at MOI = 25,000 unless otherwise indicated, not exceeding 25 % of culture volume. The next day, typically 18 h later, cells were harvested for transplantation by centrifugation, washed once in PBS and resuspend at 10^6^ cells/mL in PBS. Alternatively, cells were analyzed 2 days after electroporation unless otherwise indicated.

### Human HSPC transplantation into MISTRG6 mice

Newborn 1–3 day old pups of MISTRG6 were injected intrahepatically with 30 µL of gene-edited human fetal liver HSPC suspension (30,000 cells/mouse) using a 30-gauge needle. In some experiments pups were sublethally irradiated with 1.5 Gy just before transplantation which improved engraftment. In some experiments, also adult MISTRG6 mice were injected with 10^5^ cells whereas in another experiment some pups received human HSPCs that were cryopreserved in FBS with 10 % DMSO after gene editing and then thawed and washed as described above. Cells from at least 3 different donors were used in in vivo experiments.

### Immunizations

Mice were immunized as indicated in figures depending on the experiment. For TM4 core immunizations mice were immunized intraperitoneally with 100 µL containing 10 μg TM4 core gp140 (*56, 66*) in PBS adjuvanted with 33 % (v/v) alhydrogel (Invivogen). Sometimes animals were additionally immunized in the hind footpads with 25 uL/footpad containing 2.5 µg TM4 core gp140 in 33 % alhydrogel. For YU2 gp140 immunizations animals were immunized with 15 µg in 150 µL adjuvanted with 33 % (v/v) alhydrogel, 10 µg injected i.p and 2.5 µg into each hind footpad. For malaria antibodies, animals were immunized with a PfCSP-ferritin nanoparticle immunogen similar to the 126 immunogen previously described in (*67*). Animals received a total of 0.15 µg PfCSP-ferritin immunogen in 150 µL adjuvanted with 50 % (v/v) Sigma Adjuvant System (Sigma-Aldrich), 100 ng injected i.p and 25 ng into each hind footpad. For influenza virus experiments animals were immunized with recombinant hemagglutinin protein from strain A/Hong Kong/1/1968 (H3N2) with a C-terminal T4 bacteriophage fibritin foldon for trimerization followed by an avitag and a 6×His tag. Animals received a total of 7.5 µg in 150 µL adjuvanted with 50 % (v/v) Sigma Adjuvant System (Sigma-Aldrich), 5 µg injected i.p and 1.25 µg into each hind footpad.

Serum samples were collected throughout the experiments as indicated in the figures by submandibular bleeding and if necessary, animals were terminally bled under isoflurane anesthesia first submandibularly followed by cardiac puncture.

### B cell depletion from bone marrow and secondary chimeras

Bone marrow from primary recipients was harvested and red blood cells lysed with ACK lysing buffer as described above. Whole bone marrow cells were depleted of B cells using CD19 microbeads (Miltenyi Biotec, Cat.# 130-121-301) and LD columns (Miltenyi Biotec, Cat.# 130-042-901) according to the manufacturer’s instructions. 3 million CD19-depleted bone marrow cells/recipient in PBS were injected i.v. into lethally irradiated (2 × 5 Gy) B6.SJL secondary recipients.

### ICE analysis

Genomic DNA was extracted from single cell suspensions using QuickExtract DNA extraction solution (Biosearch Technologies, Cat#. QE09050). PCRs to amplify mouse B2M locus targeted by CRISPR/Cas9 sgRNA sgB2M, sequence CATGTGATCAAGCATCATGA or unedited controls, were performed using Phusion Green Hot Start II High-Fidelity polymerase (Thermo Fisher, Cat.# F537L) and the following primers: TCAAAGGGCATGCCAGGTAG and CAGCAGAACTGGCAGGGTTA. PCR was performed according to manufacturer’s instructions with 40 cycles, annealing at 65 °C for 30 s and extension at 72°C for 30 s. PCR product size was verified by gel electrophoresis, and bands were gel-extracted (Machery Nagel, Cat.# 740609) and sent for Sanger sequencing (Genewiz) using the indicated primers. ab1 files were analyzed using the ICE CRISPR analysis webtool (<u>ice.editco.bio</u>).

### Digital droplet PCR

Genomic DNA (gDNA) for ddPCR was extracted from single cell suspensions using a column based purification method to sufficiently fragment genomic DNA (Qiaamp DNA Micro Kit, Cat.# 56304). gDNA concentration was determined by Nanodrop reading for dsDNA (ThermoFisher) and diluted accordingly with nuclease-free water. ddPCR was performed according to manufacturer’s instructions with 100–200 ng gDNA template per reaction, primers (900 nM) and PrimeTime 6-Fluorescein amidite (FAM)- or Hexachlorofluorescein (HEX)-labelled probes (250 nM) (all IDT, see Data S5 for details) using ddPCR Supermix for Probes (No dUTP) (Bio-Rad). PCR primers to measure editing efficiency were designed to detect edited alleles by annealing one primer in the genomic region not found in HDR template and the second primer annealing to a unique sequence located in the insert with the FAM-labelled probe annealing in between. Reference PCRs used HEX-labelled probes and primers annealed on a different chromosome producing amplicons of similar size, to the matched edited allele PCR. Both assays were multiplexed in a single PCR reaction. Droplets were generated according to manufacturer’s instructions using the AutoDG system (Bio-Rad). Droplets were thermocycled on a BioRad C1000 Touch at 95 °C for 10 min; followed by 50 cycles of 94 °C for 30 s, 60 °C for 1 min and 72 °C for 2 min followed by once step of 98 °C for 10 min to stabilize droplets. Droplets were read using the QX200 Droplet reader (Bio-Rad). Data was analyzed using QX Manager Standard edition 2.0 (Bio-Rad). Percentage of edited alleles was calculated as 100×[copies/µL of edited allele]/[copies/µL of reference amplicon]. All reactions were performed and measured in technical duplicates.

### Flow cytometry

Mouse spleens were forced through a 70 µm mesh into FACS buffer (PBS containing 2 % heat-inactivated FBS and 2 mM ethylenediaminetetraacetic acid), and red blood cells were lysed in ACK lysing buffer (Gibco) for 3 min. Lymph nodes were dissociated in 1.5 mL tubes using a cell pestle and filtered through a 70 µm mesh cell strainer. Bone marrow was isolated from long bones by centrifugation as described above. Cultured cells were harvested by centrifugation. Cells were centrifuged at 200–400 g for 5–6 min at 4 °C and washed in PBS or FACS buffer. Fc-receptors were blocked for 15 min on ice. Cells were stained for 20 min on ice with primary antibodies or reagents listed in Data S6 and, depending on the stain, washed again and secondary-stained for another 20 min on ice before acquisition on a BD LSRFortessa or BD Symphony A3 or A5. All cell sorting was performed on the BD FACSymphony S6. Anti-idiotypic antibodies were produced as human IgG1/κ or mouse IgG1/κ and species mismatched with the cells for detection. Anti-idiotype binding was detected using Zenon anti-mouse IgG or anti human IgG1 reagents (PE and APC) respectively, according to manufacturer’s instructions (ThermoFisher). Biotinylated peptides for detection of anti-PfCSP antibodies and biotinylated antigens for detection of other antibodies were detected with streptavidin-fluorophore conjugates as indicated in the figures. Peptides were identical to the ones used in ELISA, details below. Data was analyzed using FlowJo 10.10.0.

### ELISAs

Costar 96-well, half area high binding polystyrene assay plates (Corning, Cat.# 3960) were coated with the indicated protein (H1 PR8, H3 HK68, TM4 core, YU2 gp140), antibody (anti-3BNC117 idiotype or anti-10-1074 idiotype as mouse or human IgG1), all produced in house, or *P. falciparum* CSP derived synthetic peptide (NPNANPNANPNANPNANPNANP-Biotin (asparagine, alanine, asparagine and proline, NANP) or NVDPNANPNVDPNANPNVDP-Biotin (asparagine, valine, aspartic acid and proline, NVDP) both from JPT Peptide Technologies) at 2 µg/mL in PBS over night at 4°C. Plates were blocked with 5 % skimmed milk powder in PBS for 2 h at room temperature. 6 washes in PBS 0.05 % Tween 20 were performed after every subsequent step. Sera were diluted in PBS at 1:50 top dilution and 1:3 (naïve and after prime) or 1:4 (after boosts) serially diluted for an 8-point curve. Recombinant antibody standards were diluted to 2 µg/mL and diluted 1:5 in PBS for an 8-point curve. Secondary antibodies conjugated to horseradish peroxidase were used to detect bound flag-tagged antibodies (1:1000, clone L5, Biolegend Cat.# 637311), StrepTag II-tagged antibodies (1:1000, IBA, Cat.# 2-1509-001) or mouse IgG (1:5000, Jackson ImmunoResearch Cat.# 115-035-071 or Southern Biotech Cat.# 1030-05) or mouse IgM or mouse IgA (Southern Biotech Cat.# 5300-05B). Binding was revealed using 3,3’,5,5’-tetramethylbenzidine substrate (ThermoFisher, Cat.# 34021) and the reaction stopped by addition of 1M H_2_SO_4_ (Sigma). Absorbance was read at 450 nm and 570 nm on a FLUOstar Omega (BMG Labtech). For analysis of monoclonal antibody concentration in serum, an identical recombinant antibody standard, if necessary with the same tag, expressed as mouse IgG1, was run on every ELISA plate and used to fit a sigmoidal 4-parameter logistic regression standard curve in GraphPad Prism 10 which was used to interpolate serum concentrations from dilutions in the exponential phase of the curve. For total anti-antigen IgG responses, sigmoidal 4-parameter logistic regression was fitted to curves of every sample and used to calculate the half maximal effective concentration (EC_50_). µg/mL values below detection were set to minimum detection level based on control sera to avoid plotting 0 on logarithmic plots. For samples were signal was too low for a curve fit EC_50_ was set to 10, 5x above the highest measured dilution.

### ELISpot

Immobilon-P membrane high protein-binding 96-well plates (Millipore, S2EM004M99) were activated with 50 µL/well of 35 % ethanol, washed 3-times with PBS and coated overnight at 4 °C with 100 µL/well of 10 µg/mL anti-3BNC117 idiotype as human IgG1/𝜅 in PBS. Plates were washed twice with PBS and blocked with B cell culture medium (see section on B cell gene editing) overnight at 4 °C. Spleen and bone marrow were harvested and red blood cells lysed as described above, then resuspended in B cell culture medium, counted and adjusted to 1.5×10^7^ live cells/mL. 3×10^6^ cells/well or another three 3-fold serial dilutions (10^6^, 0.33×10^6^ and 0.11×10^6^ cells/well) were seeded into the coated and blocked ELISpot plates and incubated at 37 °C 5 % CO_2_ for 18 h. Plates were then washed 3-times with PBS and incubated with 100 µL/well of 0.22 µm sterile-filtered anti-mouse IgG-HRP (Jackson Immunoresearch Cat# 115-035-071) for 2 h at room temperature. Plates were washed 6-times with PBS and developed with AEC substrate solution prepared to the manufacturer’s instructions (Sigma-Aldrich, Cat# AEC101) for 10 min at room temperature. Development was stopped by washing thoroughly with distilled water. Plates were left to dry for several days and imaged using Agilent BioTek Cytation 7. Spots were quantified manually.

### IgG purification from serum and HIV-1 neutralization assay

Serum IgG and HIV-1 neutralization assay were performed as previously described (*9, 68*). In brief, cryopreserved mouse serum was pooled (see Data S2) and IgG purified using protein G Ab SpinTraps (28-4083-47; GE Healthcare), then concentrated and buffer-exchanged into PBS using Amicon Ultra 30K centrifugal filter units (UFC503024; Merck Millipore) according to the manufacturer’s instructions. Neutralizing activity against a multiclade panel of 3BNC117-sensitive HIV-1 Env pseudoviruses was determined using the validated TZM.bl neutralization assay as previously described (*69*).

### *P. falciparum* sporozoite traversal inhibition assay

The sporozoite traversal inhibition assay was performed as previously described (*67*) at the Max Planck Institute for Infection Biology in accordance with local safety authorities (Landesamt für Gesundheit und Soziales Berlin, Germany, LAGeSo, project number 297/13). Briefly, *A. coluzzii* mosquitoes (*70*) were infected with mature NF54 *Plasmodium falciparum* gametocytes via artificial glass feeders for 15 min and kept at 26 °C and 80 % humidity in a controlled S3 facility. Infected mosquitoes received an additional uninfected blood meal 8 days post-infection. 60,000 human hepatocyte HC-04 cells (*71*) were seeded at 60,000 density one day before the assay in flat-bottomed 96-well plates (Corning). Sporozoites were isolated 14–15 days post-infection by grinding mosquito thoraces containing the salivary glands with glass pestles and filtered with a 40 µm cell strainer (BD Biosciences). *Pf* sporozoites (50,000) were pre-incubated for 30 min on ice in HC-04 medium with serially diluted pooled mouse serum samples (dilution range 1:25 to 1:400). The serum-sporozoite mixtures were transferred to the HC-04 cells in presence of dextran-tetramethylrhodamine (0.5 mg/mL, 10,000 MW, Molecular Probes) and incubated for 2 h at 37 °C and 5 % CO_2_. Cells were then washed, trypsinized and fixed with 1 % paraformaldehyde in PBS before flow cytometry quantification using a FACS LSR II instrument (BD Biosciences). Data analysis was performed using FlowJo 10.10.0 and for each sample traversal efficiency was calculated by subtraction of the background (dextran positive cells incubated with uninfected mosquito salivary gland material). Traversal inhibition (%) was calculated as: 100 - (traversal efficiency^SAMPLE^ / traversal efficiency^MEDIUM^ × 100).

### SHM analysis

Analysis of SHM was performed by single cell antibody sequencing according to a previously published protocol with the following modifications (*72*). Flag-tagged donor CD45.2^+^ GC B cells of mice receiving MEDI882_flag_ edited HSPCs were single cell sorted into TCL lysis buffer with 1% β-mercaptoethanol and frozen at -80°C until further processing. RNA was purified using RNAClean XP beads (Beckman Coulter) and reverse transcribed using SuperScript III (Thermo Scientific) as previously described in an 18 µL reaction but using gene specific primers for exon 1 of Ighm (CCCATGGCCACCAGATTCTT) and Ighg1, Ighg2b, Ighg2c, Ighg3 genes (AGAAGGTGTGCACACCGCTGGAC) with annealing at 55°C as per manufacturer’s instructions. 3 µL of cDNA were used as template in a 20 µL Phusion Plus (Thermo Scientific) PCR reaction according to the manufacturer’s instructions with 40 cycles of 98 °C for 30 s, 67 °C for 10 s and 72 °C for 45 s using forward primer GCAAGAGAGTGTCCGGTTAGT annealing just upstream of the translation start site and reverse primers CCCATGGCCACCAGATTCTT and AGAAGGTGTGCACACCGCTGGAC annealing in the Cµ or Cɣ constant region respectively resulting in a ∼1.3 kb fragment encompassing both the VJ_L_ and VDJ_H_ regions of edited B cells. PCR products were Sanger sequenced by Genewiz using primers GCTCAGGGAARTAGCCCTTGAC; AGGGGGAAGACATTTGGGAAGGAC; TGAGACACAGTCTTAGATATCACCAT; GGGTCAGGAGCAACCAACTT and GACACGGTGACCATAGTCCC. Sequences were analyzed manually using Geneious Prime (GraphPad). Only cells for which the entire VDJ_H_ or VJ_H_ sequences were sequenced with at least 1 read per position with a quality score higher than 30 were counted. Cells that showed mutations from the template MEDI8852_flag_ sequence were cherry-picked, reamplified and re-sequenced to exclude both PCR and sequencing errors. Additionally, the batch of recombinant MEDI8852_flag_ AAV6 used for creating the sequenced edited B cells was sequenced using Oxford Nanopore sequencing (Plasmidsaurus, Big AAV purified capsid, ITR to ITR sequencing) and no different antibody sequences other than the expected template sequence were detected.

### Influenza virus challenge

Isoflurane-anesthetized mice were intranasally infected with the indicated dose and strain of influenza virus in PBS. A/Puerto Rico/8/1934 (H1N1) was provided by the laboratory of Jeffrey Ravetch, Rockefeller University, New York USA(*73*). A/Vietnam/1203/2004 (H5N1, 2:6 PR8 reassortant with polybasic cleavage site deleted (*74*) was provided by the laboratory of Florian Krammer, Icahn School of Medicine at Mount Sinai, New York, USA. All influenza challenge experiments were performed in a BSL2+ animal facility at Rockefeller University in strict adherence to local regulations and with authorization from the Institutional Review Board and the Rockefeller University Institutional Animal Care and Use Committee.

### Recombinant protein production

Vectors used for recombinant HA trimer protein were kindly provided by laboratories of Jeffrey V. Ravetch (Rockefeller University) and Patrick C. Wilson (Weill Cornell Medical College). All constructs were confirmed by Sanger (Azenta) or Oxford Nanopore sequencing (Plasmidsaurus). Unless described otherwise below, recombinant proteins were produced in Expi293F F cells by transient transfection of expression plasmids using Expifectamine 293 transfection kit (Gibco, A14525) as previously described (*56, 66*). In brief, 6 days after transfection, culture supernatant was harvested, centrifuged to pellet cells and then filter sterilized for affinity chromatography. To purify trimeric Env proteins, supernatant was passed through an agarose-bound *Galanthus nivalis* lectin (GNL) resin (Vector Laboratories) followed by a size-exclusion column (SEC) chromatography. His-tagged proteins were purified over a Ni Sepharose 6 Fast Flow resin (Cytiva, 117531803). Peak fractions from size-exclusion chromatography were identified using native gel electrophoresis, and the fractions corresponding to monomeric Envelope or HA trimers were pooled and stored at -20 °C. Antibodies were purified over a protein G Sepharose 4Fast Flow resin (Cytive 70611805) and buffer exchanged into PBS and stored at -80°C.

Purified HA trimer protein was biotinylated using the EZ-Link-Sulfo-NHS-LC-Biotinylation kit according to the manufacturer’s instructions (Thermo Fisher Scientific, 31497). Excess biotin was removed by diafiltration with 100 kDa cutoff. Biotinylated HA was stored at -20°C.

Avi-tagged TM4 core gp120 was biotinylated using a BirA reaction according to the manufacturer’s instructions (Avidity, EC6.3.4.15), buffer exchanged to PBS and stored at -80°C.

PfCSP-ferritin immunogen containing the junction epitope, NVDP and NANP motifs, a 15-aa linker, H. pylori apoferritin, a 3-aa linker and PADRE were expressed in HEK 293F cells. Supernatants containing secreted molecules were filtered and buffer exchanged to 20 mM sodium acetate pH 4 using a tangential flow filtration LV Centramate Cassette of 100 kDa molecular weight cut-off (Pall Laboratory) and purified by ion exchange chromatography (HiTrap Q HP, Cytiva), followed by hydrophobic chromatography (HiTrap Phenyl HP, Cytiva). The protein was desalted into PBS, sterile-filtered and frozen at -80 °C.

RC1 SOSIP antigen was kindly provided by Harry B. Gristick from the laboratory of Pamela Bjorkman and produced as previously described (*75*). RC1 was randomly biotinylated as described above for HA. Excess biotin was removed by centrifugation using Amicon Ultra Centrifugal filters with a 50 kDa cutoff (Millipore). Biotinylated RC1 SOSIP was aliquoted and frozen at -80°C.

### Quantification and statistical analysis

Statistical information, including *n* and statistical significance values, are indicated in the text or the figure legends. GraphPad Prism 10 was used for statistical analysis. No statistical methods were used to predetermine sample size. The experiments were not randomized and the investigators were not blinded to allocation during experiments and outcome assessment. Age- and sex matched animals were used in between groups.

## Acknowledgements

We thank Thomas Eisenreich for help with mouse colony management and technical help; Kristie Gordon and Jean-Philip Truman for assistance with cell sorting; Masa Jankovic and Tacio A. Waldetario for laboratory support; Katherine Prieto for technical support in providing the PfCSP-ferritin immunogen; Angela Asante, Nadine Neitzel and Cornelia Kreschel for assistance with sporozoite traversal inhibition assay; Harry B. Gristick and the Bjorkman laboratory at Caltech for provision of RC1 SOSIP protein and all members of the Nussenzweig laboratory for helpful discussions.

## Author contributions

H.H. conceived the study, acquired funds, designed and performed all experiments, interpreted data, and wrote the manuscript. K.-H. Y. assisted with mouse experiments and husbandry and AAV production; C.R., P.L., G.L.D.R., P.Z., T.H., M.L, J.H., L.B. and M.J. assisted and performed experiments. T.X. and E.S. performed and R.A.F. supervised MISTRG6 experiments. B.H. and A.G. produced recombinant proteins and antibodies. M.S.S. performed and interpreted TZM-bl neutralization assays. G.C. performed and E.A.L designed and interpreted results of the sporozoite traversal inhibition assay. S.A.N., J.C., P.C.W. and F.K assisted with influenza experiments and provided reagents. S.K, J.-P. J., H.W., P.S. assisted with malaria experiments and provided reagents. L.S. provided HIV reagents. M.C.N. supervised the study, acquired funds, designed experiments, interpreted data, and wrote the manuscript. All authors edited the manuscript.

## Data and materials availability

All data needed to evaluate the conclusions in the paper are available in the main text and supplementary materials. Gene editing constructs and sequences are available from H.H. and M.C.N. under a material transfer agreement with The Rockefeller University under reasonable request.

## Supplementary Materials

Supplementary Text Figs. S1 to S16 Data S1 to S6

