## Supplementary Text and Figures for "B Lymphocyte Protein Factories produced by Hematopoietic Stem Cell Gene Editing"

**The PDF file includes:**

Supplementary text
Figs. S1 to S16

**Other Supplementary Materials for this manuscript include the following:**

Data S1 to S6

#### Supplementary Text

*Differences in antibody titers*

To give guidance to future development of this approach for different antibodies, we discuss here the observed variability in antibody titers between different antibodies and immunizations. The data indicates that the antibody paratope is the strongest determinant of the antibody titer as various antibodies differed significantly even when using identical immunizations (compare Fig. 5F-H for PfCSP-ferritin immunization with anti-malaria antibodies 317, 2514 and CIS43.D3 or Fig. 3K for YU2-gp120 immunization for anti-HIV bNAbs 3BNC117 and 10-1074).

Another important determinant is antibody affinity for the antigen. In line with published data, high affinity antigens are necessary to efficiently recruit B cells into immune response (*55, 56*). Whereas YU2 gp140, a high affinity antigen for 10-1074 induced potent 10-1074 serum responses, low affinity antigen TM4 core did not (Fig S9D). This was also seen in responses after a TM4 core boost (boost 2) in Fig. 3K for 10-1074.

Conversely, we show similar levels for 3BNC117 in TM4core immunization and YU2 gp140 immunization (compare Fig. 1F to Fig. 3K) suggesting that if antibody affinity is high for the cognate antigen, it can start a similar antibody response.

The antibody paratope will also determine whether there is any amount of polyreactivity which would also alter the half-life of the antibody. Although there is no measurable polyreactivity by either antibody, we see higher titers of 10-1074 than for 3BNC117 in the mouse models presented here (Fig. 3K) which is in line with 10-1074 having a longer half-life than 3BNC117 in humans (*58, 61*).

Lastly, autoreactivity of the human, antibody V regions we used to mouse antigens may detrimentally influence B cell development, activation and antibody expression. This may reduce precursor frequency and responsiveness to antigenic stimulation and prevent certain antibodies from efficient expression in certain animal models.

In conclusion, predicting antibody expression levels for a given antibody-immunization combination are non-obvious and will require individual testing under several different conditions but choice of non-polyreactive antibodies and use of a cognate high-affinity antigen will likely increase chances of success. Notably, the results with engineered mouse HSPCs carrying anti-HIV-1 antibodies mirror the half-lives of the antibodies in humans.

*Somatic hypermutation and adaption to evolving pathogens*

Our data indicate that somatic hypermutation is low in edited GC B cells and thus likely affinity maturation is equally low. Although we cannot exclude that in a setting of low level but chronic antigen exposure that there may be some adaptation to the virus through SHM, the likelihood seems low. However, mice are neither a good model for memory B cell responses nor HIV studies so this question will have to be addressed in non-human primates or eventually clinical trials.

While we think this method could serve as a strategy to achieve a functional cure for HIV, this would rely on the use of a combination of anti-HIV bNAbs rather than the evolution of the bNAb response. Since the vast majority of SHM is typically detrimental, silent or ineffectual, it may in fact be advantageous to have a constant source of native bNAb-expressing B cells derived from HSCs to guarantee the intended bNAb breadth and potency while also allowing some cells to affinity mature.

### Fig. S1

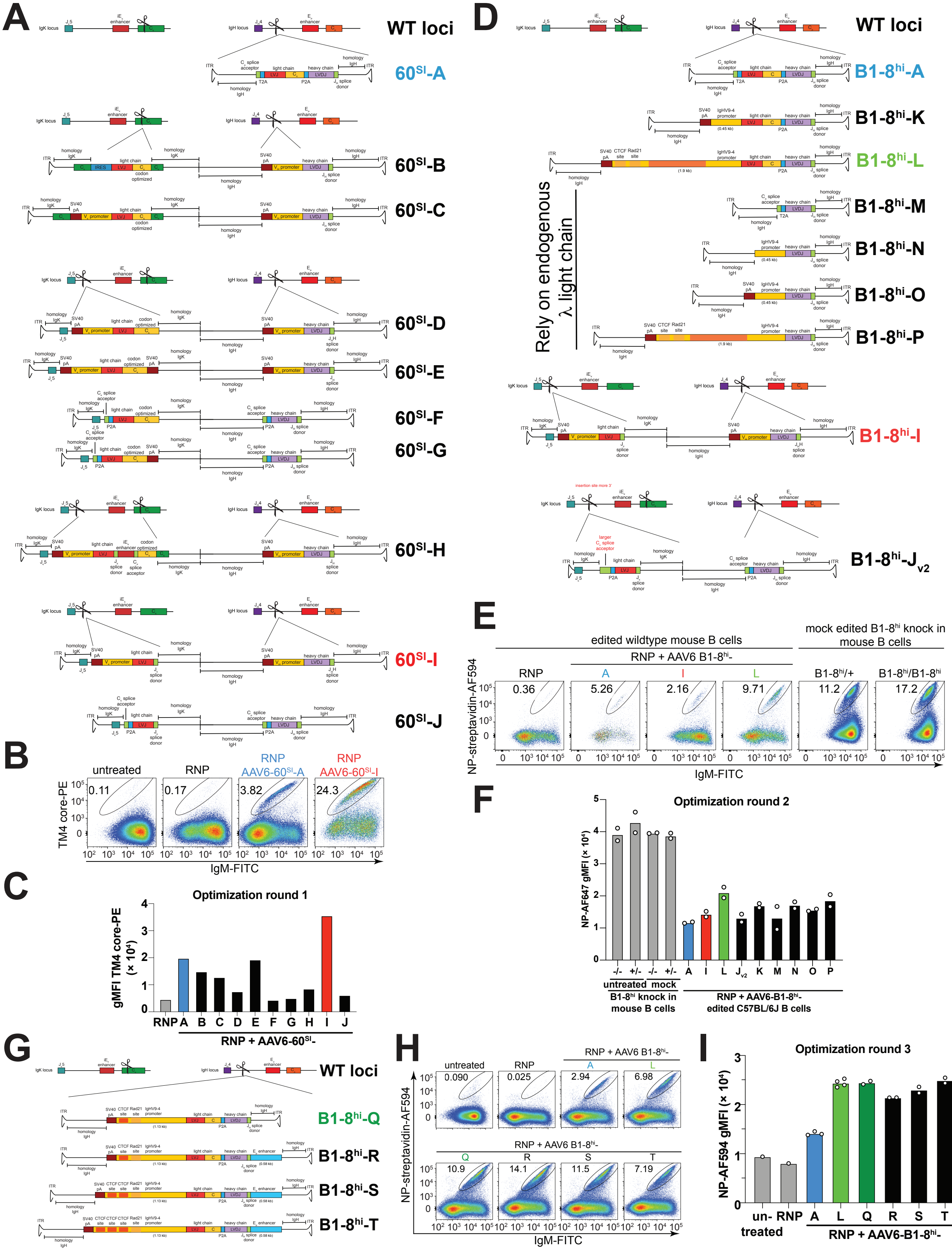

A
WT loci
60SI-A
60SI-B
60SI-C
60SI-D
60SI-E
60SI-F
60SI-G
60SI-H
60SI-I
60SI-J
B
untreated
RNP
RNP AAV6-60SI-A
RNP AAV6-60SI-I
TM4 core-PE
IgM-FITC
C
Optimization round 1
gMFI TM4 core-PE (× 10⁴)
RNP + AAV6-60SI-
D
WT loci
B1-8hi-A
B1-8hi-K
B1-8hi-L
B1-8hi-M
B1-8hi-N
B1-8hi-O
B1-8hi-P
Rely on endogenous λ light chain
B1-8hi-I
B1-8hi-Jv2
E
edited wildtype mouse B cells
mock edited B1-8hi knock in mouse B cells
RNP + AAV6 B1-8hi-
NP-streptavidin-AF594
IgM-FITC
F
Optimization round 2
NP-AF647 gMFI (× 10⁴)
untreated mock
B1-8hi knock in mouse B cells
RNP + AAV6-B1-8hi- edited C57BL/6J B cells
G
WT loci
B1-8hi-Q
B1-8hi-R
B1-8hi-S
B1-8hi-T
H
untreated
RNP
RNP + AAV6 B1-8hi-
NP-streptavidin-AF594
IgM-FITC
I
Optimization round 3
NP-AF594 gMFI (× 10⁴)
un-treated
RNP + AAV6-B1-8hi-

**Fig. S1. Optimization of BCR expression in gene-edited mouse B cells.** Related to Fig. 1. 3 rounds of optimization of constructs for BCR expression were performed. Benchmark construct A highlighted in blue, initial lead constructs I highlighted in red, optimized constructs L and Q highlighted in light and dark green respectively throughout the figure. Construct L and Q were used for subsequent experiments. **(A)** Schematic of anti-HIV-1 antibody 3BNC60$^{SI}$ (60$^{SI}$) constructs and insertions in the mouse *IgH* and/or *IgK* locus testing separate targeting of light and heavy chains into their respective loci. All constructs were packaged as AAV6 and transduced into activated primary mouse B cells after Cas9 RNP electroporation using sgRNAs cutting at the indicated locations (scissors). L, Leader; V, V gene; D, D gene and J, J gene. (**B**) Exemplary flow cytometric analysis of 3BNC60$^{SI}$ expression by TM4 core antigen binding 2 days after gene editing as in (A). (**C**) Quantification of (A, B). Geometric mean fluorescence intensity of TM4 core on edited cells gated as in (B). (A-C) Representative of 2 independent experiments. (**D**) Schematic of second round of constructs based on results from (A-C) using NP-specific antibody B1-8$^{hi}$ which can pair with endogenous λ light chains to test expression of constructs encoding only a B1-8$^{hi}$ heavy chain (B1-8$^{hi}$-M, -N, O, -P) or introducing a longer promoter containing CTCF- and Rad21-binding sites for transcriptional insulation (B1-8$^{hi}$-L, -P) or an improved version of construct J ($J_{v2}$). Constructs were benchmarked against B1-8$^{hi}$ versions of previous benchmark construct A (blue) and lead construct I (red) as well as untreated or mock electroporated B cells from B1-8$^{hi}$ heterozygous (+/-) or homozygous (-/-) knock in mice. (**E**) Exemplary flow cytometric analysis of B1-8$^{hi}$ expression by NP-antigen binding 2 days after gene editing as in (D). (**F**) Quantification of (D, E). Geometric mean fluorescence intensity of antigen NP-streptavidin-AF647 on edited cells gated as in (E). Each dot represents a technical replicate. (D-F) Representative of 2 independent experiments. (**G**) Schematic of third generation constructs based on results from (D-F). Based on the latest lead construct B1-8$^{hi}$-L (light green) containing a long promoter with insulator elements, a shorter construct containing an internal deletion from the Rad21 binding site to the original core promoter was tested (construct Q, dark green). Additionally, the $E_{\mu}$ enhancer was added to some constructs and the CTCF-binding site was dupli- or triplicated. (**H**) Flow cytometric analysis of benchmark and third generation constructs for B1-8$^{hi}$ expression by NP-antigen binding 2 days after gene editing as in (G). (**I**) Geometric mean fluorescence intensity of antigen NP-streptavidin-AF594 on edited cells gated as in (H). Each dots represents a technical replicate. Numbers in flow cytometric plots indicate percentage of cells within gate of all cells plotted.

### Fig. S2

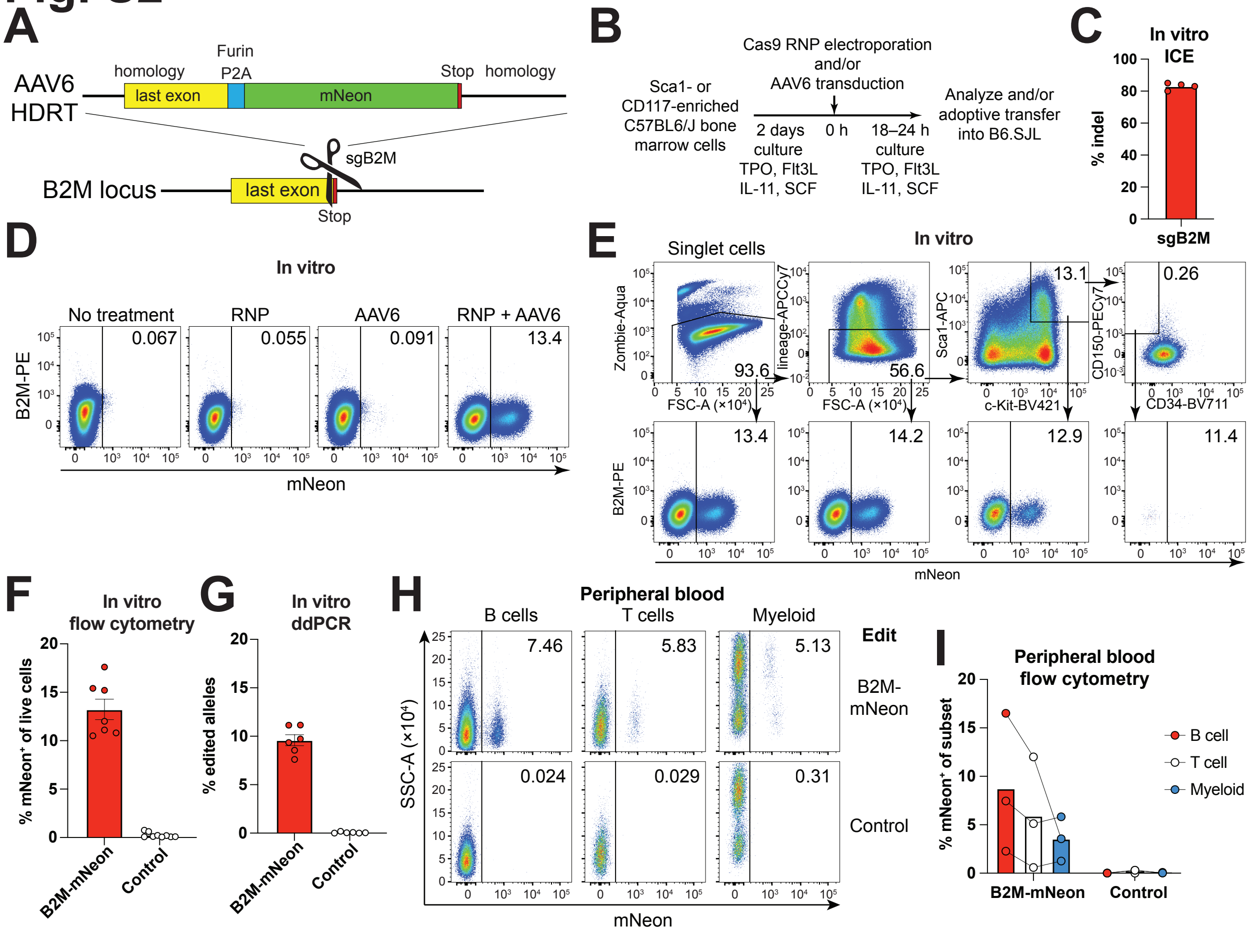
A
AAV6
HDRT
homology
last exon
Furin
P2A
mNeon
Stop
homology
sgB2M
B2M locus
last exon
Stop
B
Sca1- or
CD117-enriched
C57BL6/J bone
marrow cells
Cas9 RNP electroporation
and/or
AAV6 transduction
2 days
culture
TPO, Flt3L
IL-11, SCF
0 h
18–24 h
culture
TPO, Flt3L
IL-11, SCF
Analyze and/or
adoptive transfer
into B6.SJL
C
In vitro
ICE
% indel
sgB2M
D
In vitro
No treatment
RNP
AAV6
RNP + AAV6
0.067
0.055
0.091
13.4
B2M-PE
mNeon
E
In vitro
Singlet cells
Zombie-Aqua
FSC-A (×10⁴)
93.6
lineage-APCCy7
56.6
Sca1-APC
c-Kit-BV421
13.1
CD150-PECy7
CD34-BV711
0.26
B2M-PE
13.4
14.2
12.9
11.4
mNeon
F
In vitro
flow cytometry
% mNeon⁺ of live cells
B2M-mNeon
Control
G
In vitro
ddPCR
% edited alleles
B2M-mNeon
Control
H
Peripheral blood
B cells
T cells
Myeloid
Edit
7.46
5.83
5.13
B2M-
mNeon
0.024
0.029
0.31
Control
SSC-A (×10⁴)
mNeon
I
Peripheral blood
flow cytometry
% mNeon⁺ of subset
B cell
T cell
Myeloid
B2M-mNeon
Control

**Fig. S2. Gene editing of mouse HSPCs.** Related to Fig. 1 **(A)** Gene editing strategy to tag the mouse B2M gene with mNeon. The locus is targeted with an sgRNA close to the B2M stop codon (stop). Recombinant AAV6 is packaged with a repair template tagging the B2M CDS with a furin-cleavage site, a glycine-serine-glycine (GSG)-linker, and a P2A self-cleaving oligopeptide followed by the mNeon gene. The insert is flanked by 821 bp (5') and 811 bp (3') homology arms. **(B)** Experimental layout for mouse HSPC gene editing. Bone marrow from wildtype C57BL/6J mice is enriched for Sca1 expressing cells by magnetic positive selection. Cells are put into culture in serum free medium with the indicated cytokines for 2 days. Then cells are electroporated with Cas9 RNPs and subsequently transduced with the recombinant AAV6 HDRT. 18–24 h after transfection cells are analyzed or adoptively transferred into congenically marked B6.SJL mice. (**C**) Cutting efficiency of Cas9 at the B2M locus in mouse B cells using sgB2M analyzed using gDNA target locus PCR amplification, PCR product Sanger sequencing and ICE algorithm analysis of the chromatogram (**D**). Expression of mNeon by flow cytometry in cultured mouse HSPCs 2 days after electroporation after the indicated treatment. Gated on live, singlet, cells. (**E**) Data as in D but subgated into HSC phenotype cells. (**F**) Quantification of (D) Each dot represents a technical replicate. Pool of 4 independent experiments. (**G**) ddPCR measuring correct target integration at the B2M locus using in-out primer strategy on genomic DNA from bulk cultures 1 day after electroporation. Each dot represents a technical replicate of 3 biological replicates from 1 experiment (**H**) Flow cytometric analysis of mNeon expression in donor $CD45.2^{+}$ leukocyte subsets 6 weeks after transplantation of B2M-mNeon or irrelevantly-edited control HSPCs. (**I**) Quantification of (H). Each dot represents an individual mouse from 1 independent experiment. Numbers in flow cytometric plots indicate percentage of cells within gate of all cells plotted.

#### Fig. S3

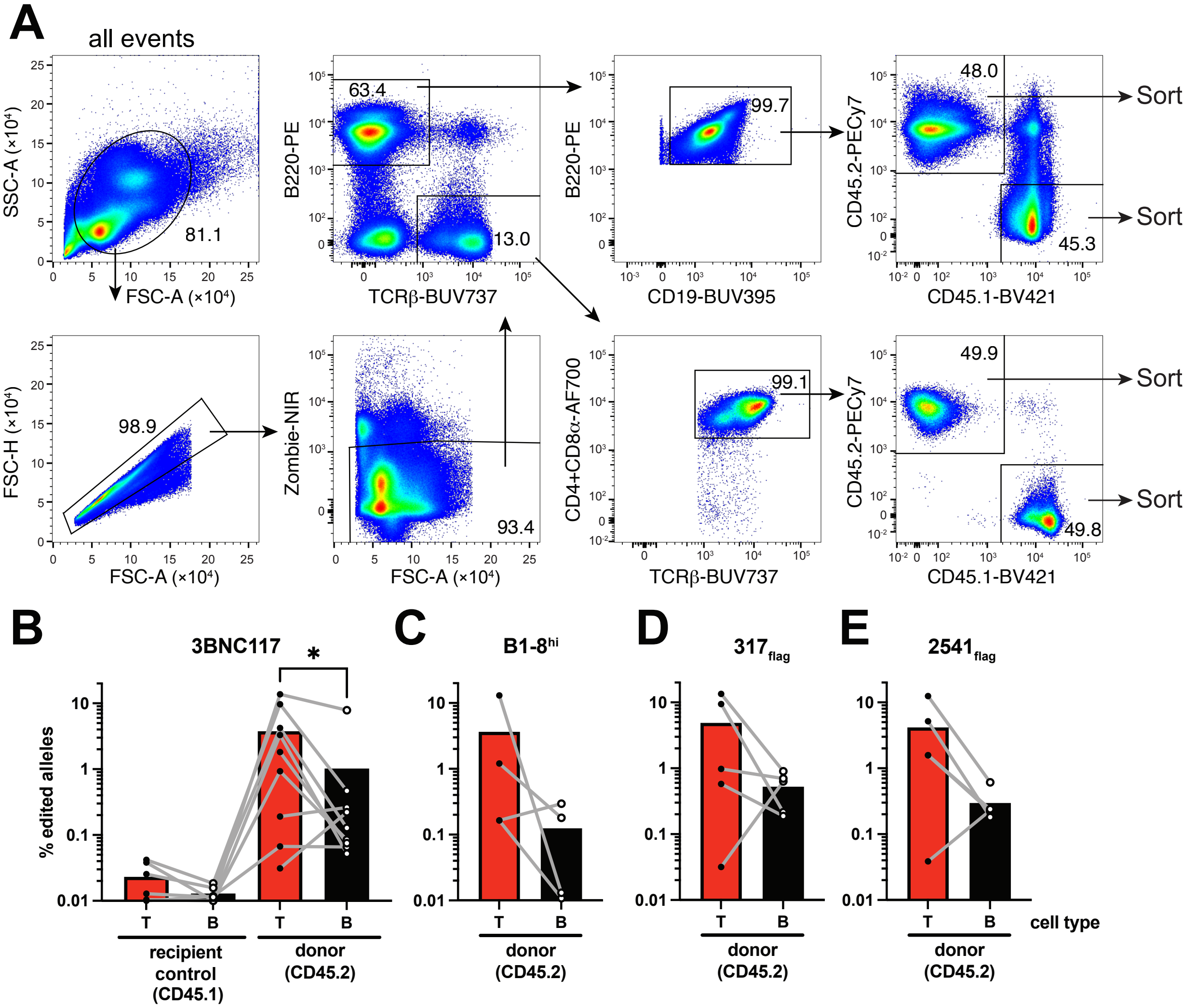

A
all events
SSC-A (×10⁴)
FSC-A (×10⁴)
81.1
FSC-H (×10⁴)
98.9
Zombie-NIR
93.4
B220-PE
TCRβ-BUV737
63.4
13.0
CD19-BUV395
99.7
CD4+CD8α-AF700
99.1
CD45.2-PECy7
CD45.1-BV421
48.0
45.3
49.9
49.8
Sort
B
3BNC117
% edited alleles
recipient control (CD45.1)
donor (CD45.2)
C
B1-8hi
D
317flag
E
2541flag
T
B
cell type

**Fig. S3. Gene editing-efficiency in B and T cells *in vivo*.** Related to Fig. 1 **(A)** Gating strategy for B cell and T cell purification for ddPCR from mouse peripheral blood shown in (B)-(E)**.** Numbers in plots indicate percentage of cells within gate of all cells plotted. **(B)** Gene-editing efficiency *in vivo* in recipient control ($CD45.1^+$) or donor ($CD45.2^+$) B ($CD19^+$ $B220^+$) or T ($TCR\beta^+$ $CD4/CD8\alpha^+$) cells from peripheral blood of mice receiving 3BNC117-edited HSPCs 6-10 weeks after reconstitution by digital droplet PCR measuring target locus integration using an in-out primer strategy. Pooled data from 2-3 independent experiments measured in 1–2 technical replicates is shown, each dot representing an individual mouse, paired data from the same mouse connected by a grey line. Bars indicate mean. B cell data points are also part of Fig 1D. (**C**) As in (B) but mice reconstituted with B1-$8^{hi}$ or (**D**) $317_{flag}$ or (**E**) $2541_{flag}$ edited HSPCs. 0 values set to 0.01 for display on logarithmic axis.* indicated $p = 0.03$ in a paired T test.

### Fig. S4

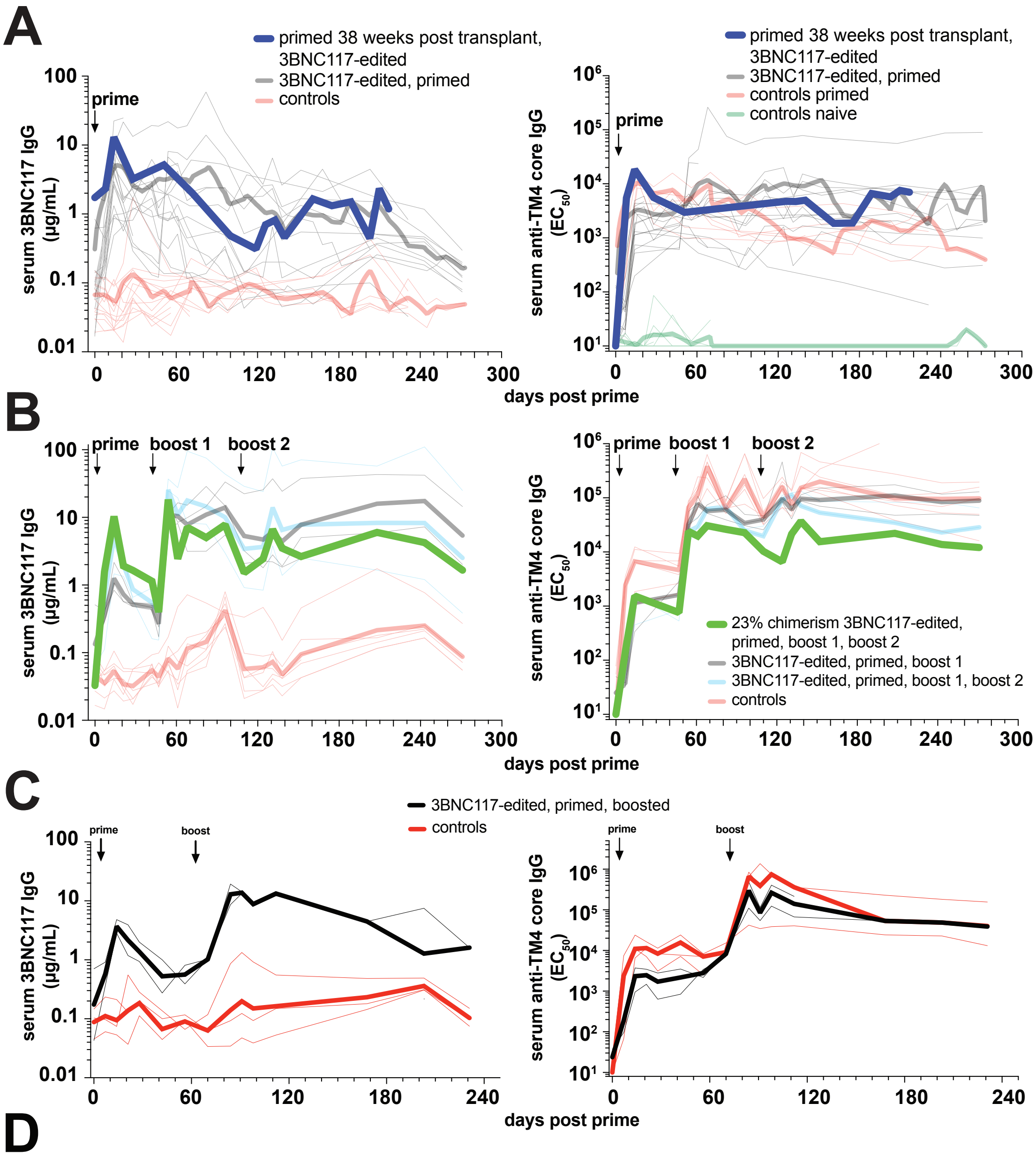


# D

| | Polyclonal purified IgG $IC_{50}$ Titer in TZM.bl cells (μg/mL) | | | | | | | | | | | | ELISA | | |
|---|---|---|---|---|---|---|---|---|---|---|---|---|---|---|---|
| | Q23.17 (Clade A) | | RHPA4259.7 (Clade B) | | Q769.d22 (Clade C) | | 001428-2.42 (Clade C) | | 263-8 (CRF02_AG) | | 3BNC117-resistant control Du422.1 (Clade C) | | anti-idiotypic 3BNC117 IgG | | HSPC |
| Sample ID | $IC_{50}$ | $IC_{80}$ | $IC_{50}$ | $IC_{80}$ | $IC_{50}$ | $IC_{80}$ | $IC_{50}$ | $IC_{80}$ | $IC_{50}$ | $IC_{80}$ | $IC_{50}$ | $IC_{80}$ | (μg/mL) | TM4 core immunizations | gene-edit |
| ckit-5 | >184 | >184 | >184 | >184 | >184 | >184 | >184 | >184 | >184 | >184 | >184 | >184 | 14.418 | prime | 3BNC117 |
| Sca1-8 | >132 | >132 | >132 | >132 | >132 | >132 | >132 | >132 | >132 | >132 | >132 | >132 | 1.420 | prime | 3BNC117 |
| Sca3-12 | **34.328** | >106 | **17.334** | **50.933** | **15.721** | **64.220** | **13.784** | **41.369** | **47.485** | >106 | >106 | >106 | 33.048 | prime | 3BNC117 |
| Sca2-4 | **49.744** | >114 | **22.959** | **73.091** | **18.959** | **66.328** | **37.714** | **101.496** | **64.015** | >114 | >114 | >114 | 32.476 | prime, boost day 47 | 3BNC117 |
| Sca3-8 | **8.898** | **53.680** | **4.642** | **16.498** | **5.221** | **20.582** | **4.723** | **18.176** | **20.703** | **80.429** | >105 | >105 | 105.438 | prime, boost day 47 | 3BNC117 |
| Sca3-20 | **58.671** | >126 | **22.994** | **74.215** | **14.602** | **98.497** | **24.053** | **85.884** | **75.502** | >126 | >126 | >126 | 25.573 | prime, boost day 47 | 3BNC117 |
| Sca4-9 | >109 | >109 | >109 | >109 | >109 | >109 | >109 | >109 | >109 | >109 | >109 | >109 | 4.253 | prime, boost day 70 | 3BNC117 |
| Sca5-3 | **62.135** | **147.219** | **20.927** | **76.584** | **28.486** | **86.201** | **17.264** | **51.617** | **72.346** | >176 | >176 | >176 | 24.182 | prime, boost day 70 | 3BNC117 |
| Sca5-12 | >98 | >98 | **31.139** | >98 | **46.692** | >98 | >98 | >98 | >98 | >98 | >98 | >98 | 20.782 | prime, boost day 70 | 3BNC117 |
| Sca4-18 | >178 | >178 | >178 | >178 | >178 | >178 | >178 | >178 | >178 | >178 | >178 | >178 | 3.48 | neg. ctr, prime, boost day 70 | B1-8$^{hi}$ |
| Sca3-21 | >137 | >137 | >137 | >137 | >137 | >137 | >137 | >137 | >137 | >137 | >137 | >137 | 1.074 | neg. ctr, prime, boost day 47 | none |
| Sca3-22 | >118 | >118 | >118 | >118 | >118 | >118 | >118 | >118 | >118 | >118 | >118 | >118 | 0.510 | neg. ctr, prime, boost day 47 | none |
| Sca3-24 | >106 | >106 | >106 | >106 | >106 | >106 | >106 | >106 | >106 | >106 | >106 | >106 | 2.188 | neg. ctr, prime, boost day 47 | none |
| Sca3-25 | >126 | >126 | >126 | >126 | >126 | >126 | >126 | >126 | >126 | >126 | >126 | >126 | 2.988 | neg. ctr, prime, boost day 47 | none |
| Sca3-26 | >137 | >137 | >137 | >137 | >137 | >137 | >137 | >137 | >137 | >137 | >137 | >137 | 3.031 | neg. ctr, prime, boost day 47 | none |
| Mouse 3BNC117 IgG1 | **0.026** | **0.073** | **0.022** | **0.060** | **0.022** | **0.078** | **0.009** | **0.034** | **0.039** | **0.151** | >1.0 | >1.0 | N/A | N/A - Recombinant antibody | N/A |
| Human 3BNC117-LS-J | **0.008** | **0.036** | **0.008** | **0.025** | **0.009** | **0.041** | **0.003** | **0.011** | **0.012** | **0.054** | >1.0 | >1.0 | N/A | N/A - Recombinant antibody | N/A |

**Fig. S4. Detailed analysis of mice receiving 3BNC117-edited HSPCs.** Related to Fig. 1. **(A)** Serum 3BNC117 IgG and anti-TM4 core IgG data from Fig. 1E highlighting a mouse primed 38 weeks after transplant. See Data S1, mouse 2kit-5 for details. (**B)** As above but data from Fig. 1F highlighting a mouse with 23 % chimerism. (**C**) Experimental set up and analysis as in Fig. 1F but mice were boosted 70 days after prime. (**D**) TZM.bl HIV neutralization assay and anti-3BNC117 idiotype IgG ELISA of protein G purified, polyclonal mouse IgG from pooled serum of 4 to 6 time points of indicated mice with the indicated treatment >7 days after last immunization or at least 54 days after prime (sample IDs Sca2-X, Sca3-X day 54-96 pool; Sca4-X, Sca5-X days 84-112 pool, Sca1-8 days 203-259 pool; ckit-5 days 161-217 pool). Each sample corresponds to 1 mouse or recombinant antibody. Mice from 6 independent experiments are shown. Du422.1, a strain resistant to 3BNC117 and unrelated to the immunizing antigen (426c derived TM4 core) served as negative control (neg. ctr.) for unspecific neutralization activity. Irrelevantly edited B1-$8^{hi}$ or unedited wildtype mice served as experimental neg. ctr. All neg. ctr. in gray. Recombinant, monoclonal 3BNC117 expressed as mouse IgG1 or human IgG LS-J were used as positive controls. See Data S2 for details on serum pooling.

### Fig. S5

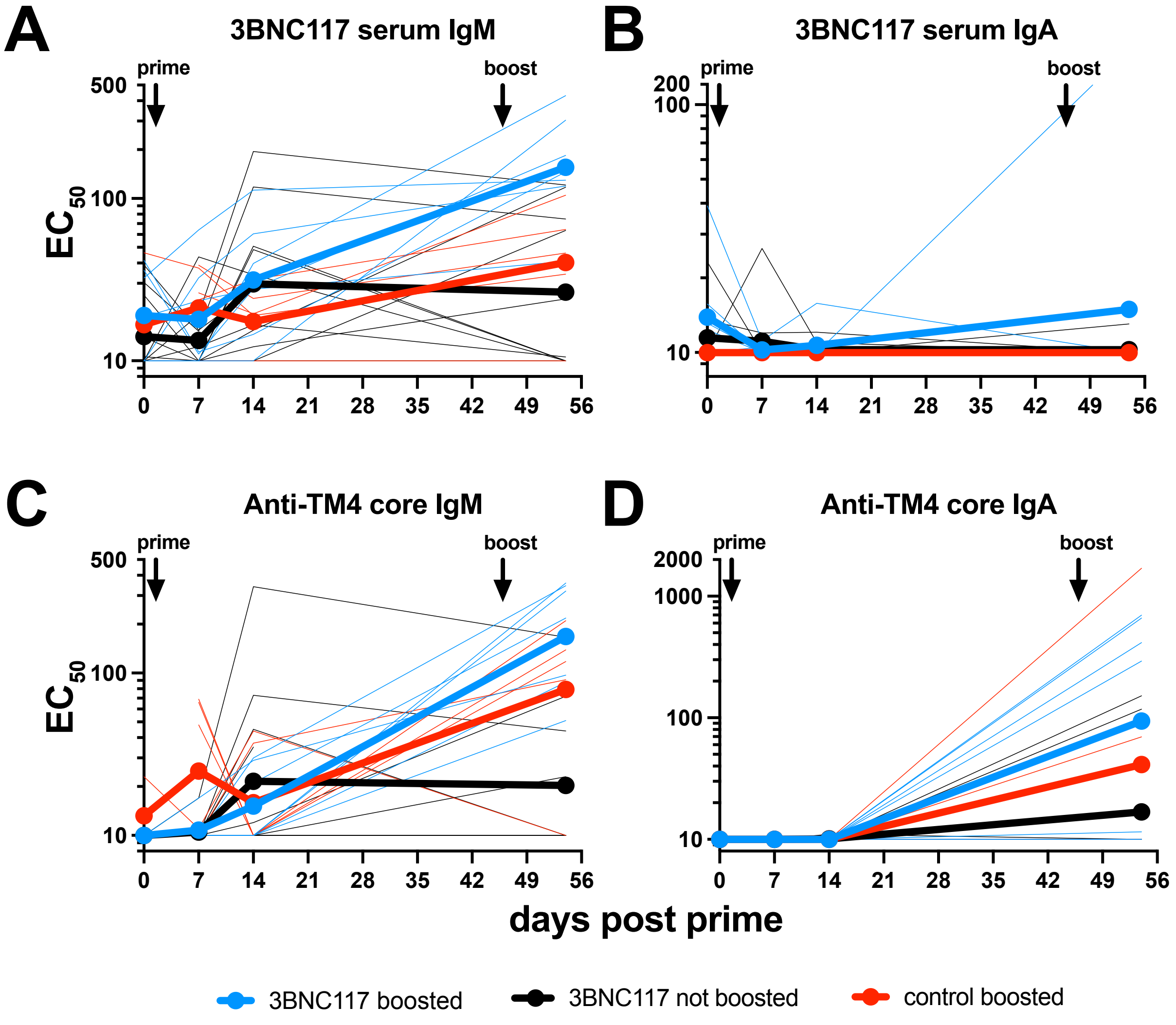
A
3BNC117 serum IgM
B
3BNC117 serum IgA
C
Anti-TM4 core IgM
D
Anti-TM4 core IgA
prime
boost
EC50
500
100
10
200
2000
1000
0
7
14
21
28
35
42
49
56
days post prime
3BNC117 boosted
3BNC117 not boosted
control boosted

**Fig. S5. IgM and IgA responses in TM4 core immunized 3BNC117-edited mice.** Related to Fig. 1 (**A**) 3BNC117 IgM or (**B**) 3BNC117 IgA in serum at the indicated time points measured by anti-idiotypic antibody ELISA and detection of the indicated antibody isotype in mice from Fig 1F. (**C**) Polyclonal anti-TM4 core IgM or (**D**) anti-TM4 core IgA in the same mice.

### Fig. S6

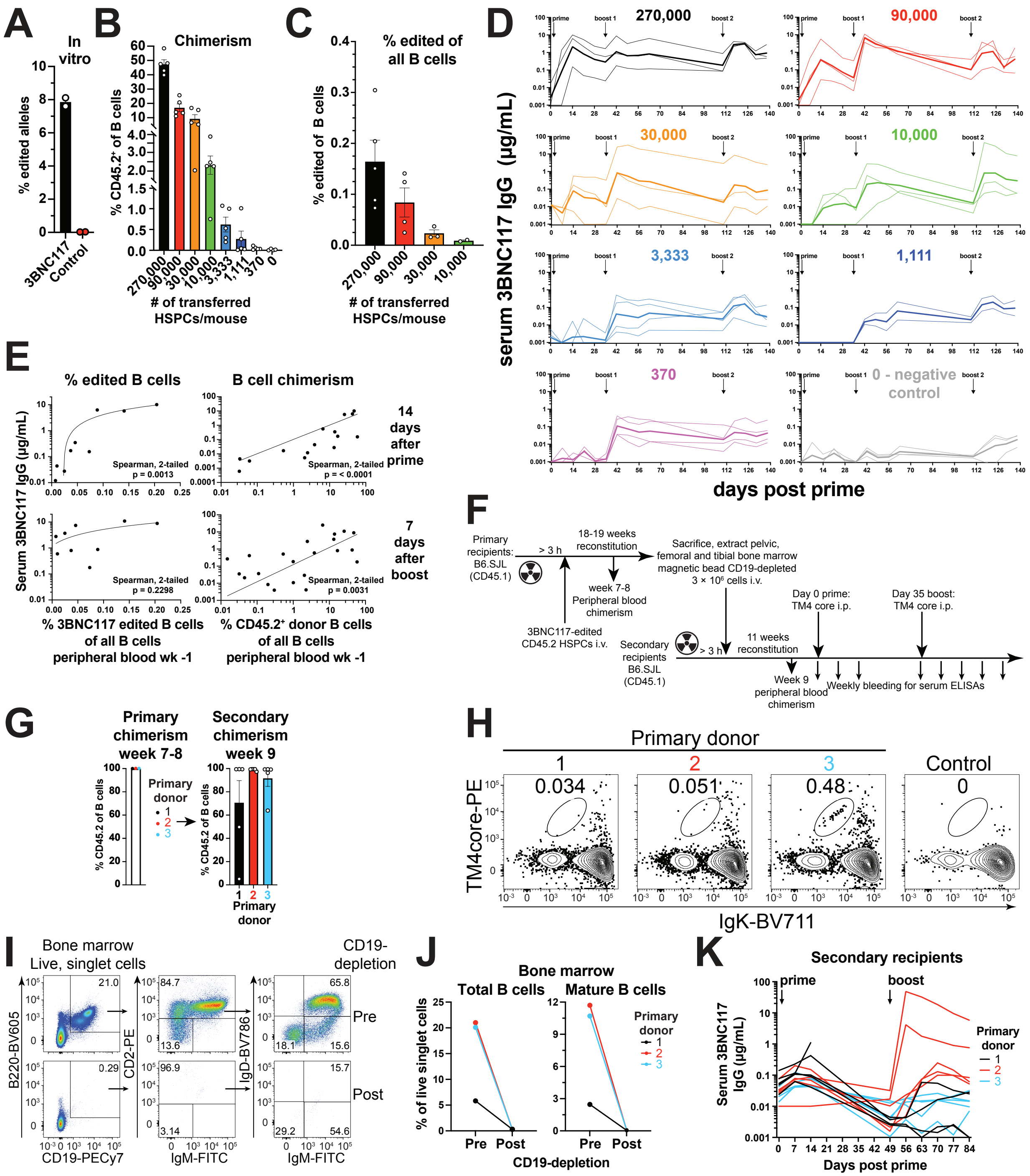
A
In vitro
% edited alleles
3BNC117
Control
B
Chimerism
% CD45.2+ of B cells
### of transferred HSPCs/mouse
C
% edited of all B cells
% edited of B cells
### of transferred HSPCs/mouse
D
270,000
90,000
30,000
10,000
3,333
1,111
370
0 - negative control
prime
boost 1
boost 2
serum 3BNC117 IgG (µg/mL)
days post prime
E
% edited B cells
B cell chimerism
14 days after prime
7 days after boost
Serum 3BNC117 IgG (µg/mL)
Spearman, 2-tailed p = 0.0013
Spearman, 2-tailed p = < 0.0001
Spearman, 2-tailed p = 0.2298
Spearman, 2-tailed p = 0.0031
% 3BNC117 edited B cells of all B cells peripheral blood wk -1
% CD45.2+ donor B cells of all B cells peripheral blood wk -1
F
Primary recipients: B6.SJL (CD45.1)
> 3 h
18-19 weeks reconstitution
week 7-8 Peripheral blood chimerism
3BNC117-edited CD45.2 HSPCs i.v.
Sacrifice, extract pelvic, femoral and tibial bone marrow magnetic bead CD19-depleted 3 × 10^6 cells i.v.
Secondary recipients B6.SJL (CD45.1)
> 3 h
11 weeks reconstitution
Week 9 peripheral blood chimerism
Day 0 prime: TM4 core i.p.
Day 35 boost: TM4 core i.p.
Weekly bleeding for serum ELISAs
G
Primary chimerism week 7-8
Secondary chimerism week 9
% CD45.2 of B cells
Primary donor
1
2
3
H
Primary donor
1
2
3
Control
0.034
0.051
0.48
0
TM4core-PE
IgK-BV711
I
Bone marrow Live, singlet cells
CD19-depletion
B220-BV605
CD19-PECy7
CD2-PE
IgM-FITC
IgD-BV786
Pre
Post
21.0
0.29
84.7
13.6
96.9
3.14
65.8
18.1
15.6
15.7
29.2
54.6
J
Bone marrow
Total B cells
Mature B cells
% of live singlet cells
Primary donor
Pre
Post
CD19-depletion
K
Secondary recipients
prime
boost
Serum 3BNC117 IgG (µg/mL)
Primary donor
Days post prime

**Fig. S6. Mixed bone marrow chimera edited HSPC titration and secondary recipients.** Related to Fig. 2 **(A)** ddPCR measuring 3BNC117 antibody editing in cultured HSPCs before transfer in the experiment depicted in Fig. 2A. Dots indicate 2 technical replicates, bars indicate mean. Unedited cells were used as control. (**B**) Chimerism and **(C)** Percentage of 3BNC117-edited B cells in peripheral blood 6 weeks after transplant and 1–2 days before prime in groups from Fig. 2A-C. Cells were gated and sorted as in Fig. S3A and antibody editing was measured in CD45.2$^+$ B cells by ddPCR. Groups with fewer transplanted HSPCs than 10,000 were below the limit of detection by ddPCR. Bars indicate mean ± SEM. Each dot indicates an individual mouse. (**D**) 3BNC117 IgG serum levels by anti-3BNC117 idiotype IgG ELISA in individual groups of mixed bone marrow chimeras from Fig. 2B. Thick lines indicate geometric mean. Thin lines indicate individual mice. (**E**) Correlations of percentage of edited B cells in the blood from (C) (left) or donor B cell chimerism from (B) (right) and 3BNC117 IgG serum levels 14 days after prime (top) or 7 days after boost (bottom). Lines indicate linear regressions. Each dot indicates a mouse. Two-tailed Spearman correlations are indicated for each graph. (**F**) Schematic experimental set up of the secondary chimera experiment shown in Fig. 2D. 1 or 2 primary donors from 2 independent experiments were used to generate 5 secondary recipients per primary donor. (**G**) Chimerism in primary (left) and secondary recipients (right). Determined by flow cytometry gated as in Fig S3A. (**H**) Flow cytometric analysis of antigen TM4 core binding on CD45.2$^+$ B cells in primary recipients 8–9 weeks after transplantation. Wildtype C57BL/6J blood was used as negative control. (**I**) Flow cytometric analysis of the B cell compartment in secondary donor bone marrow from primary recipients before and after magnetic bead depletion of CD19$^+$ cells and before transplantation into secondary recipients. (**J**) Quantification of (I). Total B cells gated as CD19$^+$ B220$^+$. Mature B cells gated as CD19$^+$ B220$^+$, CD2$^{+,}$ IgM$^+$, IgD$^+$. Each dot indicates one mouse and lines connect paired samples. (**K**) Longitudinal 3BNC117 serum levels by anti-3BNC117 idiotype IgG ELISA in secondary recipients treated as in (E), as summarized in Fig. 2D. Numbers in flow cytometric plots indicate percentage of cells within gate of all cells plotted.

### Fig. S7

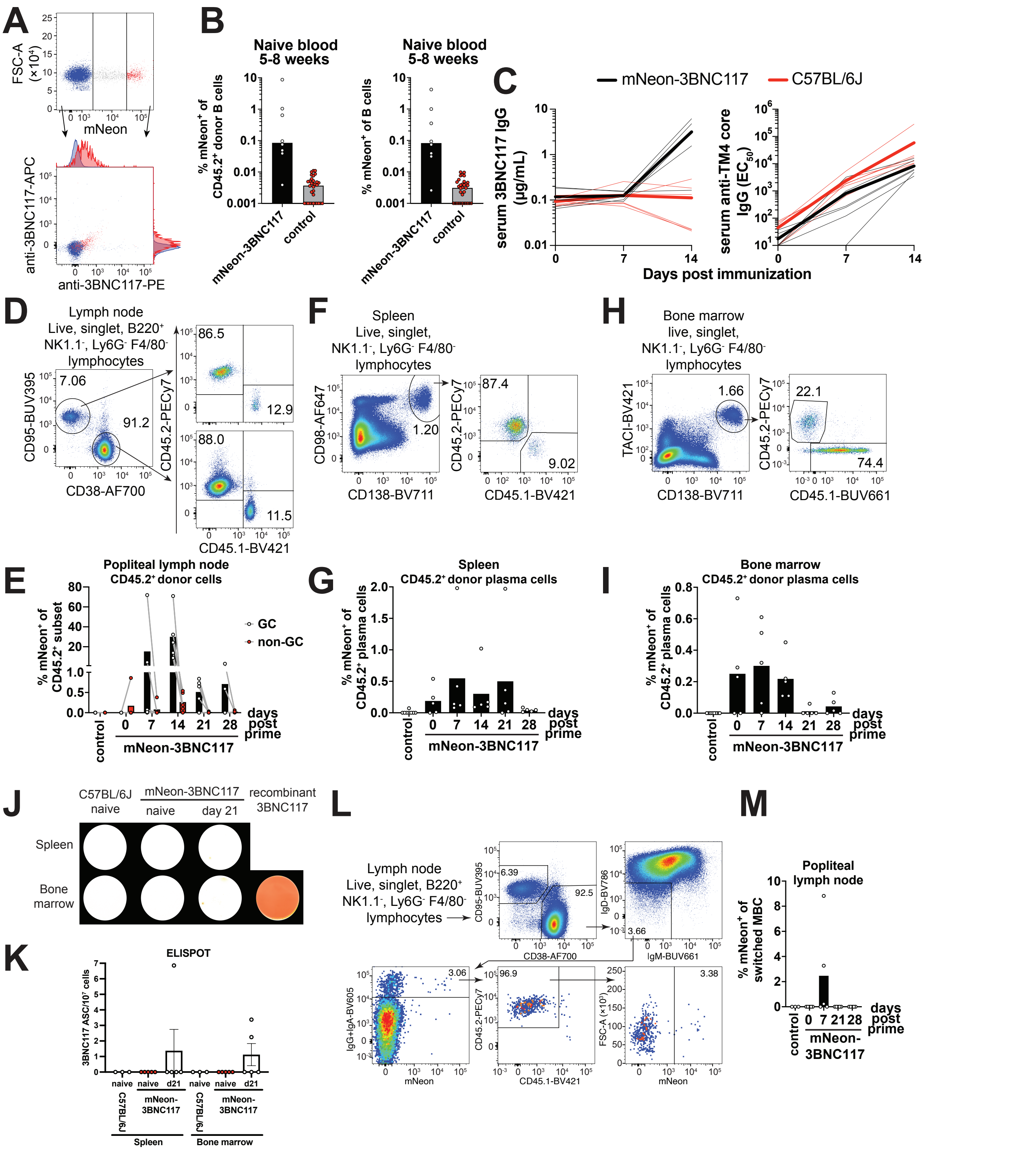
A
FSC-A (×10⁴)
mNeon
anti-3BNC117-APC
anti-3BNC117-PE
B
Naive blood 5-8 weeks
% mNeon⁺ of CD45.2⁺ donor B cells
% mNeon⁺ of B cells
mNeon-3BNC117
control
C
mNeon-3BNC117
C57BL/6J
serum 3BNC117 IgG (μg/mL)
serum anti-TM4 core IgG (EC50)
Days post immunization
D
Lymph node
Live, singlet, B220⁺
NK1.1⁻, Ly6G⁻ F4/80⁻
lymphocytes
CD95-BUV395
CD38-AF700
CD45.2-PECy7
CD45.1-BV421
7.06
91.2
86.5
12.9
88.0
11.5
F
Spleen
Live, singlet,
NK1.1⁻, Ly6G⁻ F4/80⁻
lymphocytes
CD98-AF647
CD138-BV711
1.20
87.4
9.02
H
Bone marrow
live, singlet,
NK1.1⁻, Ly6G⁻ F4/80⁻
lymphocytes
TACI-BV421
1.66
22.1
74.4
CD45.1-BUV661
E
Popliteal lymph node
CD45.2⁺ donor cells
% mNeon⁺ of CD45.2⁺ subset
GC
non-GC
days post prime
G
Spleen
CD45.2⁺ donor plasma cells
% mNeon⁺ of CD45.2⁺ plasma cells
I
Bone marrow
CD45.2⁺ donor plasma cells
J
C57BL/6J naive
mNeon-3BNC117
naive
day 21
recombinant 3BNC117
Spleen
Bone marrow
K
ELISPOT
3BNC117 ASC/10⁷ cells
naive
d21
Spleen
Bone marrow
L
Lymph node
Live, singlet, B220⁺
NK1.1⁻, Ly6G⁻ F4/80⁻
lymphocytes
6.39
92.5
IgD-BV786
IgM-BUV661
3.66
IgG+IgA-BV605
3.06
96.9
FSC-A (×10³)
3.38
M
Popliteal lymph node
% mNeon⁺ of switched MBC

**Fig. S7. Co-expression of a cargo protein, and differentiation upon immunization.** Related to Fig. 3. **(A)** Flow cytometry of peripheral blood B cells 6 weeks after transplantation, showing correlation of mNeon and 3BNC117 expression edited as in Fig. 3A, B. (**B**). Quantification of mNeon$^+$ cells among CD45.2$^+$ donor (left) or all (right) B cells in peripheral blood 6–8 weeks after transplantation. Each dot represents a mouse; bars indicate median; pool of 2 independent experiments. Controls contain untreated mice and mice receiving irrelevantly edited mouse HSPCs. (**C**) anti-3BNC117 IgG (left) and anti-TM4 core IgG ELISAs of immunized C57BL/6J controls or mice receiving gene-edited mNeon-3BNC117 mouse HSPC. Thin lines indicate individual mice and thick line indicates mean. Pool of 2 independent experiments. (**D**) Germinal center and chimerism flow cytometric stain of popliteal lymph nodes 14 days after immunization. (**E**) Quantification of mNeon-3BNC117 B cells in popliteal lymph nodes over 28 days after cognate antigen immunization among the indicated donor CD45.2$^+$ subsets gated as in (D). Each dot represents a mouse; bars indicate mean, connecting lines indicate data from the same mouse. Pool of 4 independent experiments with 2 experiments per time point. (**F**) Flow cytometric stain of splenic plasma cells and chimerism 14 days after immunization. (**G**) Quantification of mNeon-3BNC117 plasma cells in spleen over 28 days after cognate antigen immunization among donor CD45.2$^+$ plasma cells gated as in (F). Each dot represents a mouse; bars indicate median. Pool of 4 independent experiments with 2 experiments per time point. (**H**) Flow cytometric stain of bone marrow plasma cells 14 days after immunization. (**I**) Quantification of mNeon-3BNC117 plasma cells and chimerism in bone marrow over 28 days after cognate antigen immunization among donor CD45.2$^+$ plasma cells gated as in (H). Each dot represents a mouse; bars indicate median. Pool of 4 independent experiments with 2 experiments per time point. (**J**) Anti-3BNC117 idiotype specific IgG ELISpot assay of bone marrow cells or splenocytes, naïve or 21 days after TM4 core prime immunization of mice receiving mNeon-3BNC117-edited HSPCs or naïve C57BL/6J control animals. (**K**) Quantification of antibody secreting cells (ASCs) in (J). Each dots represents a mouse, bars indicate mean ± SEM. (**L**) Flow cytometric gating strategy for detection of mNeon-3BNC117-expressing isotype-switch memory B cells of draining lymph node 7 days after TM4 core immunization from mice reconstituted with mNeon-3BNC117 HSPCs. To detect all IgG$^+$ and IgA$^+$ switched cells, antibodies to IgG1a, IgG2a, IgG2b, IgG3 and IgA were added with the same fluorophore. (**M**) Quantification of (L) over 28 days. Each dot represents a mouse; bars indicate mean. Numbers in flow cytometric plots indicate percentage of cells within gate of all cells plotted.

### Fig. S8

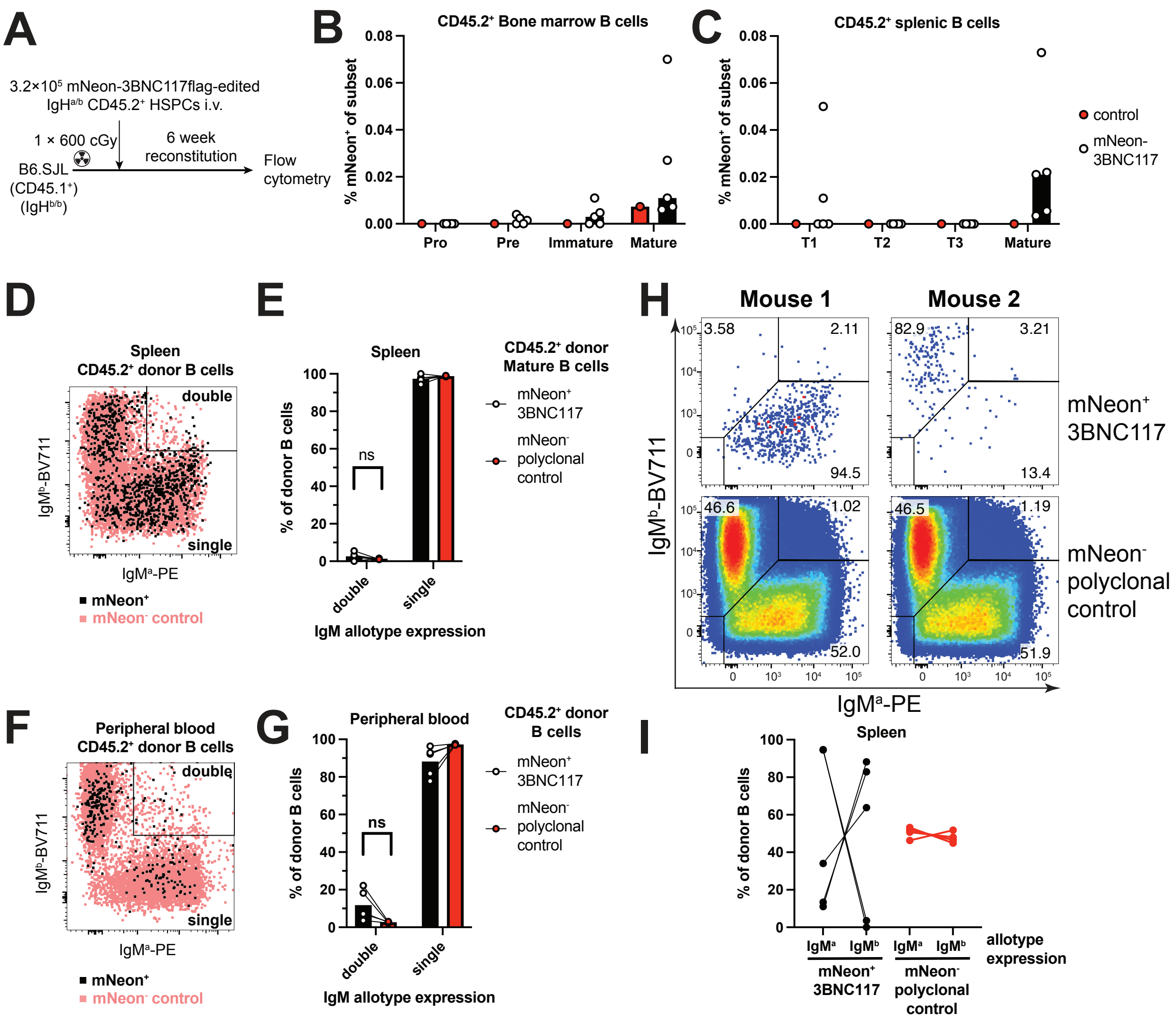

A
3.2×10⁵ mNeon-3BNC117flag-edited IgHa/b CD45.2+ HSPCs i.v.
1 × 600 cGy
6 week reconstitution
B6.SJL (CD45.1+) (IgHb/b)
Flow cytometry
B
CD45.2+ Bone marrow B cells
% mNeon+ of subset
Pro
Pre
Immature
Mature
C
CD45.2+ splenic B cells
% mNeon+ of subset
T1
T2
T3
Mature
control
mNeon-3BNC117
D
Spleen CD45.2+ donor B cells
double
single
IgMb-BV711
IgMa-PE
mNeon+
mNeon- control
E
Spleen
CD45.2+ donor Mature B cells
mNeon+ 3BNC117
mNeon- polyclonal control
% of donor B cells
ns
double
single
IgM allotype expression
H
Mouse 1
Mouse 2
3.58
2.11
94.5
82.9
3.21
13.4
46.6
1.02
52.0
46.5
1.19
51.9
mNeon+ 3BNC117
mNeon- polyclonal control
IgMb-BV711
IgMa-PE
F
Peripheral blood CD45.2+ donor B cells
double
single
IgMb-BV711
IgMa-PE
mNeon+
mNeon- control
G
Peripheral blood
CD45.2+ donor B cells
mNeon+ 3BNC117
mNeon- polyclonal control
% of donor B cells
ns
double
single
IgM allotype expression
I
Spleen
% of donor B cells
IgMa
IgMb
IgMa
IgMb
allotype expression
mNeon+ 3BNC117
mNeon- polyclonal control

**Fig. S8 Zygosity of gene-editing and heavy chain allelic exclusion.** (**A**) Schematic of the experiment presented in this figure. Allotypically marked ($CD45.2^+$) $IgH^{a/b}$ heterozygous donor HSPCs were gene-edited with the mNeon-3BNC117$_{flag}$ construct and used to reconstitute $CD45.1^+$ recipient mice. After reconstitution blood (6 weeks after transfer) or bone marrow and spleen (10 weeks after transfer) were analyzed for $mNeon^+$ cells in the B cell compartment and usage of $IgH^a$ or $IgH^b$ allele evaluated by flow cytometry. (**B**) Contribution of edited mNeon-3BNC117 B lineage cells to the indicated donor ($CD45.2^+$) bone marrow compartments measured by flow cytometry. Gating similar to Figure S6I. All compartments $CD19^+$ $B220^+$: Pro (Pro B cell, $CD2^-$); Pre (Pre B cell; $CD2^+$, $IgM^{a-}$ $IgM^{b-}$); Immature (Immature B cells, $CD2^+$, $IgM^{a+}$ or $IgM^{b+}$, $IgD^-$); Mature (Mature recirculating B cells, $CD2^+$, $IgM^{a+}$ or $IgM^{b+}$, $IgD^+$). Control is C57BL/6J. Each dot represents one mouse, bars indicate mean. (**C**) Contribution of edited mNeon-3BNC117 B lineage cells to the indicated donor $CD45.2^+$ splenic B cell compartments measured by flow cytometry. All compartments $CD19^+$ $B220^+$: T1 (transitional type 1 B cell, $CD93^+$, $CD23^-$); T2 (transitional type 2 B cell, $CD93^+$, $CD23^+$, $IgM^{a\text{-}hi}$ or $IgM^{b\text{-}hi}$); T3 (transitional type 3 B cell, $CD93^+$, $CD23^+$, $IgM^{a\text{-}low}$ or $IgM^{b\text{-}low}$); Mature (Mature B cell, $CD93^-$). Control is C57BL/6J. Each dot represents one mouse, bars indicate mean. (**D**) Flow cytometric overlay plot showing $IgM^{a+}$ $IgM^{b+}$ double expressing B cells among polyclonal $mNeon^-$ control or $mNeon^+$ 3BNC117 B cells in the spleen. Concatenate of 5 mice. (**E**) Quantification of (D) showing double ($IgM^{a+}$ $IgM^{b+}$) or single ($IgM^{a+}$ or $IgM^{b+}$) expressing B cells among polyclonal $mNeon^-$ control or $mNeon^+$ 3BNC117 B cells in the spleen. Bars indicate mean, paired data from one mouse for the same population connected by a line. Each dot indicates an individual mouse; ns indicates $p>0.05$ in paired T test. (**F**) As in (D) but in peripheral blood. (**G**) As in (E) but in peripheral blood. (**H**) Example flow cytometric plot of IgM allele bias in polyclonal $mNeon^-$ control or $mNeon^+$ 3BNC117 splenic B cells. Numbers in plots indicate percentage of cells within gate of all cells plotted (**I**) Quantification of (H) showing percentage of $IgM^{a+}$ or $IgM^{b+}$ single expressing B cells among polyclonal $mNeon^-$ control or $mNeon^+$ 3BNC117 B cells in the spleen. Paired data from the same mouse connected by a line. Each dot indicates an individual mouse.

### Fig. S9

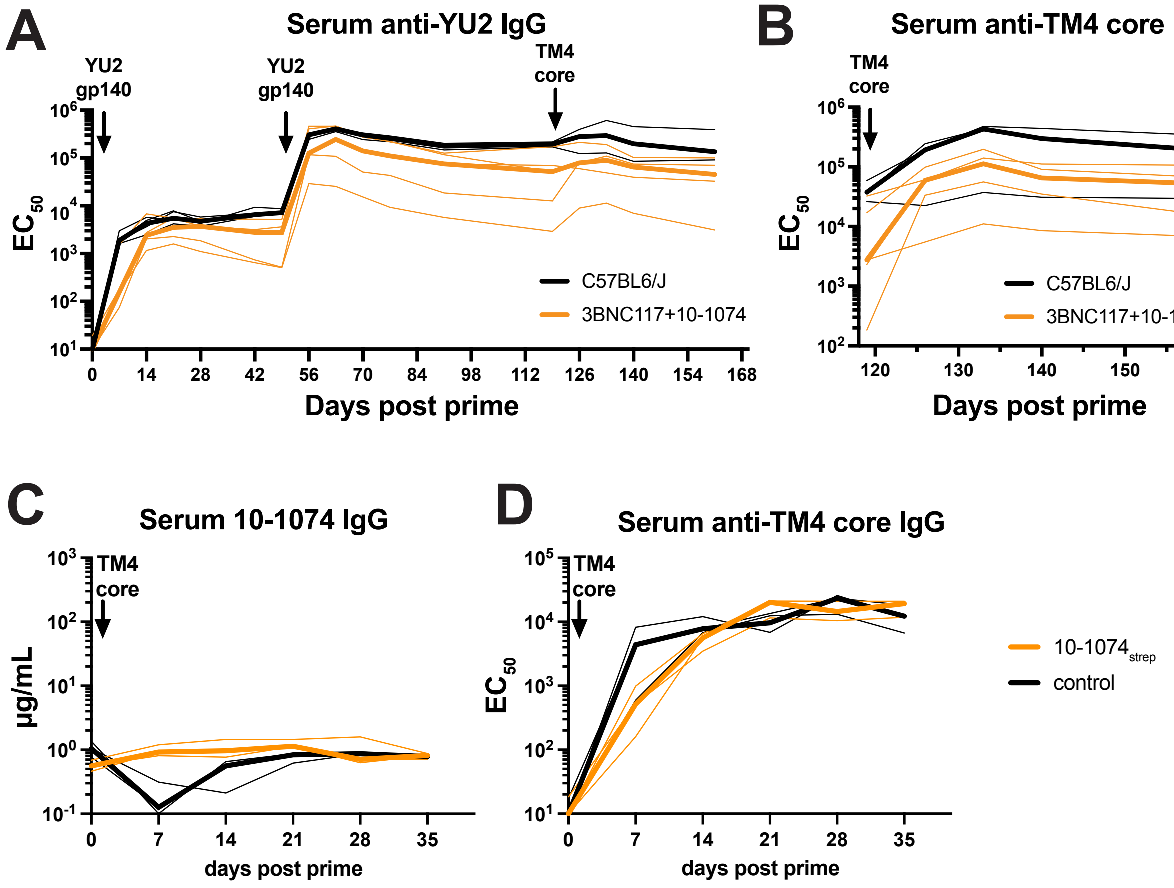
A
Serum anti-YU2 IgG
YU2 gp140
YU2 gp140
TM4 core
EC50
C57BL6/J
3BNC117+10-1074
Days post prime
B
Serum anti-TM4 core IgG
TM4 core
EC50
C57BL6/J
3BNC117+10-1074
Days post prime
C
Serum 10-1074 IgG
TM4 core
µg/mL
days post prime
D
Serum anti-TM4 core IgG
TM4 core
EC50
days post prime
10-1074strep
control

**Fig. S9. Combination of two antibodies. Related to Figure 3** (**A**) anti-YU2 gp140 and (**B**) anti-TM4 core IgG serum levels by ELISAs in mice receiving a mix of separately edited 3BNC117 and 10-1074 edited HSPCs or unedited control animals after reconstitution and YU2-gp140 prime and boost and TM4 core secondary boost immunization. Representative of 2 independent experiments. Thin lines indicate individual mice, thick lines indicate a group's median. (**C**) Serum 10-1074 IgG levels after immunization with low affinity antigen TM4 core in mice reconstituted with 10-$1074_{strep}$ edited HSPCs. (**D**) as in (C) but measuring anti-TM4 core IgG.

#### Fig. S10

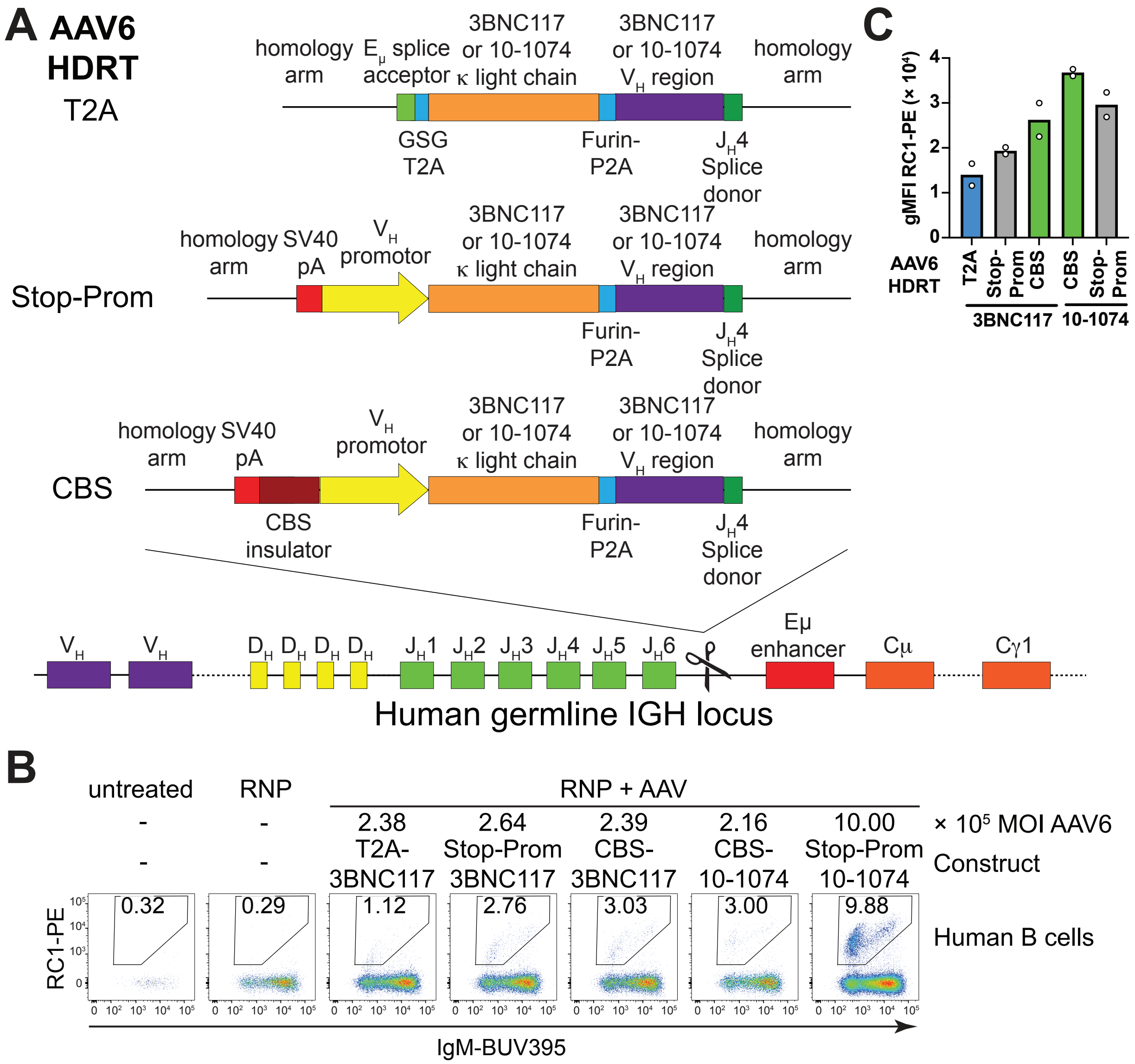

A AAV6 HDRT
T2A
homology arm
Eμ splice acceptor
3BNC117 or 10-1074 κ light chain
3BNC117 or 10-1074 VH region
homology arm
GSG T2A
Furin-P2A
JH4 Splice donor
Stop-Prom
homology arm
SV40 pA
VH promotor
CBS
CBS insulator
VH
DH
JH1
JH2
JH3
JH4
JH5
JH6
Eμ enhancer
Cμ
Cγ1
Human germline IGH locus
B
untreated
RNP
RNP + AAV
2.38
2.64
2.39
2.16
10.00
× 10⁵ MOI AAV6
T2A-3BNC117
Stop-Prom 3BNC117
CBS-3BNC117
CBS-10-1074
Stop-Prom 10-1074
Construct
0.32
0.29
1.12
2.76
3.03
3.00
9.88
Human B cells
RC1-PE
IgM-BUV395
C
gMFI RC1-PE (× 10⁴)
AAV6 HDRT
T2A
Stop-Prom
CBS
3BNC117
10-1074

**Fig. S10. Optimization of BCR expression in gene edited human B cells.** Related to Fig. 4. **(A)** Schematic of 3BNC117 or 10-1074 constructs and insertions in the human *IgH* locus testing a promoterless (T2A), promoter driven (Stop-Prom) or CTCF-binding site (CBS)-containing, promoter-driven construct design. All constructs were packaged as AAV6 and transduced into activated primary human B cells after Cas9 RNP electroporation using sgRNAs cutting at the indicated locations (scissors). (**B**) Flow cytometric analysis of bNAb expression by RC1 antigen binding 2 days after gene editing. Representative of 2 independent experiments using 2 different donors. Numbers in plots indicate percentage of cells within gate of all cells plotted (**C**) Geometric mean fluorescence intensity of antigen RC1-binding by edited B cells gated as in (B). Bar color indicates equivalent constructs to the mouse constructs of the same color in Figure S1. Bars indicate mean, dots represent technical replicates. Representative of 2 independent experiments using 2 different donors.

Fig. S11

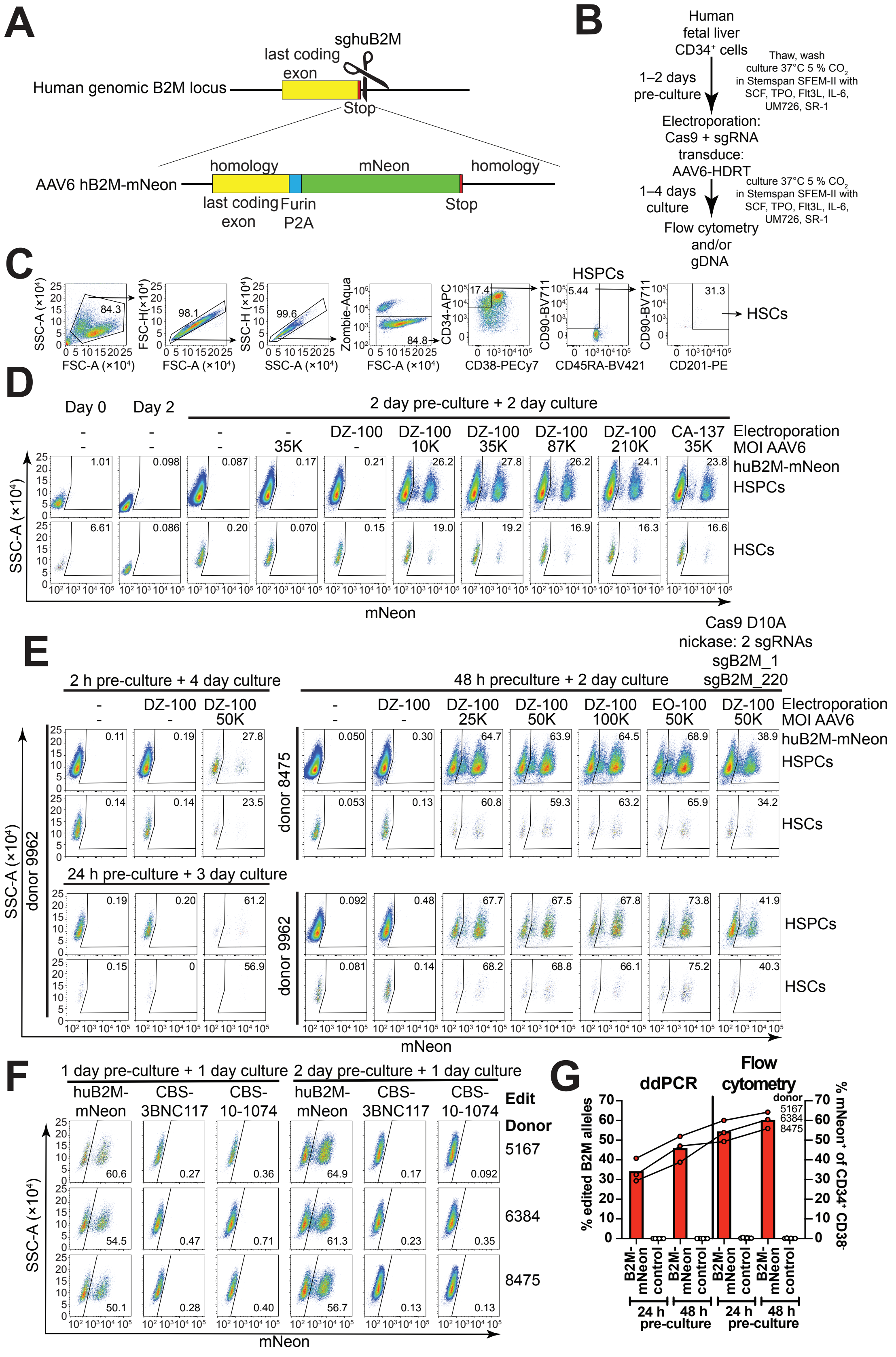
A
Human genomic B2M locus
last coding exon
sghuB2M
Stop
AAV6 hB2M-mNeon
homology
mNeon
homology
last coding exon
Furin P2A
Stop
B
Human fetal liver CD34⁺ cells
Thaw, wash culture 37°C 5 % CO2 in Stemspan SFEM-II with SCF, TPO, Flt3L, IL-6, UM726, SR-1
1–2 days pre-culture
Electroporation: Cas9 + sgRNA transduce: AAV6-HDRT
1–4 days culture
culture 37°C 5 % CO2 in Stemspan SFEM-II with SCF, TPO, Flt3L, IL-6, UM726, SR-1
Flow cytometry and/or gDNA
C
SSC-A (×10⁴)
FSC-A (×10⁴)
FSC-H (×10⁴)
SSC-H (×10⁴)
SSC-A (×10⁴)
Zombie-Aqua
CD34-APC
CD38-PECy7
HSPCs
CD90-BV711
CD45RA-BV421
CD201-PE
HSCs
84.3
98.1
99.6
84.8
17.4
5.44
31.3
D
Day 0
Day 2
2 day pre-culture + 2 day culture
DZ-100
CA-137
35K
10K
87K
210K
Electroporation
MOI AAV6
huB2M-mNeon
HSPCs
HSCs
SSC-A (×10⁴)
mNeon
E
Cas9 D10A nickase: 2 sgRNAs sgB2M_1 sgB2M_220
2 h pre-culture + 4 day culture
48 h preculture + 2 day culture
24 h pre-culture + 3 day culture
DZ-100
EO-100
25K
50K
100K
donor 9962
donor 8475
SSC-A (×10⁴)
mNeon
F
1 day pre-culture + 1 day culture
2 day pre-culture + 1 day culture
huB2M-mNeon
CBS-3BNC117
CBS-10-1074
Edit
Donor
5167
6384
8475
SSC-A (×10⁴)
mNeon
G
ddPCR
Flow cytometry
% edited B2M alleles
% mNeon⁺ of CD34⁺ CD38⁻
donor
B2M-mNeon
control
24 h
48 h
pre-culture

**Fig. S11. Optimization of human CD34$^+$ fetal liver gene editing.** Related to Fig. 4. (**A**) Gene editing strategy to tag the human B2M gene with mNeon. The locus is targeted with an sgRNA close to the B2M stop codon (stop). Recombinant AAV6 is packaged with a repair template tagging the B2M CDS with a furin-cleavage site, a glycine-serine-glycine (GSG)-linker, and a P2A self-cleaving oligopeptide followed by the mNeon coding sequence. The insert is flanked by 929 bp (5') and 903 bp (3') homology arms. (**B**) Experimental set up for (C-G). Culturing times were optimized and are indicated for each figure. (**C**) Full gating strategy for (D-F) and Fig. 4A.(**D**) Flow cytometric analysis of mNeon expression in primitive HSPCs (HSPCs, CD34$^+$, CD38$^-$) and LT-HSC phenotype cells (HSCs, CD34$^+$, CD38$^-$, CD90$^+$, CD45RA$^-$, CD201$^+$) after 0-4 days in culture using the indicated electroporation settings and multiplicity of infection for the AAV6 packaged repair template illustrated in (A). (**E**) As in (D) but varying pre-culture times and using multiple donors. (**F**) As in (E) but with different culture length and donors. Gated on primitive HSPCs (CD34$^+$, CD38$^-$) (**G**) Quantification of (F). Percentage of mNeon expressing cells among primitive HSPCs (CD34$^+$, CD38$^-$) by flow cytometry (right y axis), and percentage of edited alleles in the same samples by ddPCR (left y-axis). Irrelevantly edited cells were used as control. Connected bars indicate the same donor/sample. Bars indicate mean. Numbers in flow cytometric plots indicate percentage of cells within gate of all cells plotted

### Fig. S12

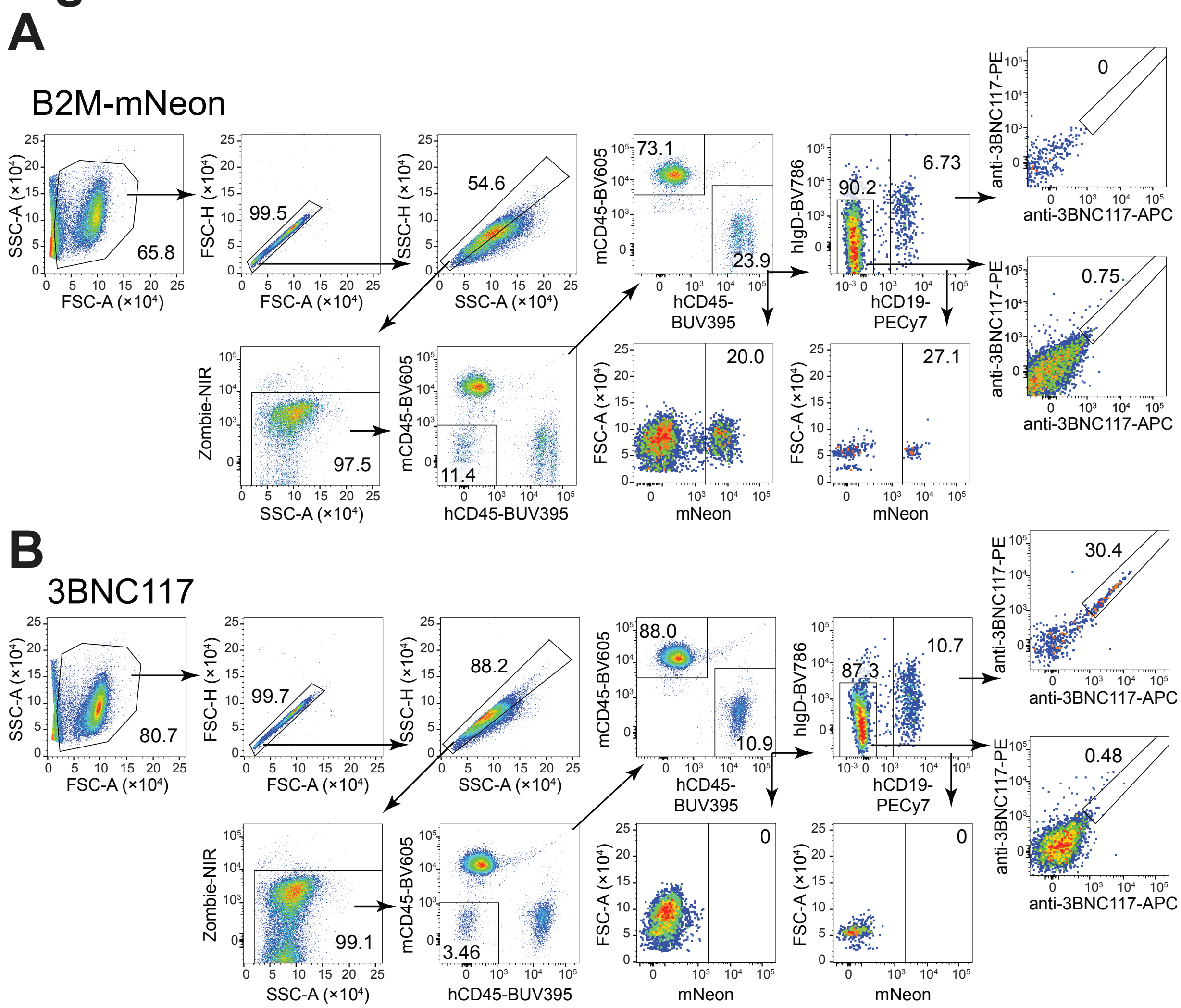
A
B2M-mNeon
SSC-A (×10⁴)
FSC-A (×10⁴)
65.8
FSC-H (×10⁴)
99.5
SSC-H (×10⁴)
SSC-A (×10⁴)
54.6
mCD45-BV605
hCD45-BUV395
73.1
23.9
hIgD-BV786
hCD19-PECy7
90.2
6.73
anti-3BNC117-PE
anti-3BNC117-APC
0
0.75
Zombie-NIR
97.5
11.4
FSC-A (×10⁴)
mNeon
20.0
27.1
B
3BNC117
80.7
99.7
88.2
88.0
10.9
87.3
10.7
30.4
0.48
99.1
3.46
0
0

**Fig. S12. Gating strategy for peripheral blood from MISTRG6 mice.** Related to Fig. 4. Full gating strategy for peripheral blood of MISTRG6 mice receiving either (**A**) B2M-mNeon or (**B**) 3BNC117-edited human $CD34^+$ fetal liver cells. Numbers in flow cytometric plots indicate percentage of cells within gate of all cells plotted

#### Fig. S13

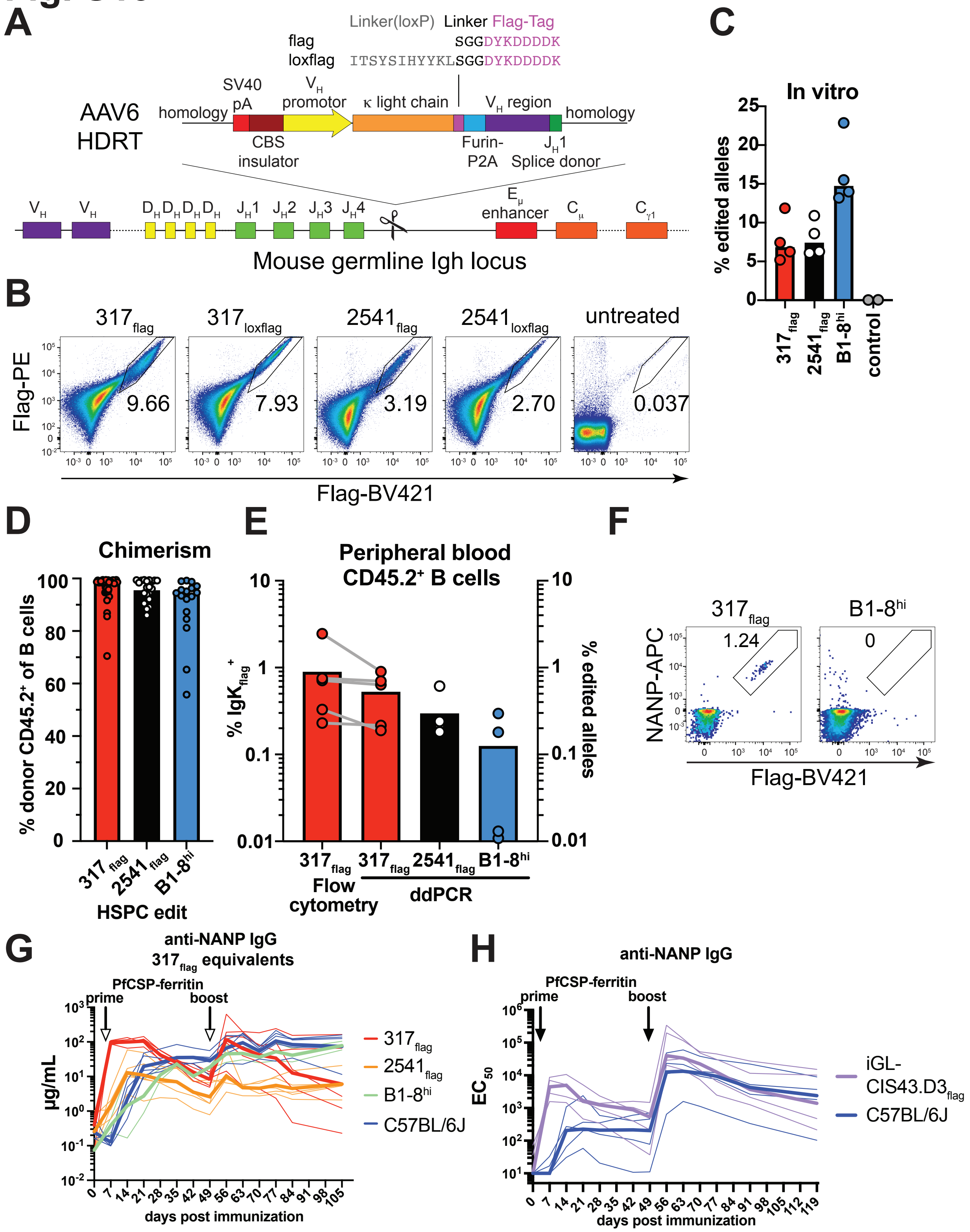

A
Linker(loxP) Linker Flag-Tag
flag SGGDYKDDDDK
loxflag ITSYSIHYYKLSGGDYKDDDDK
AAV6 HDRT
homology
SV40 pA
CBS insulator
VH promotor
κ light chain
Furin-P2A
VH region
JH1 Splice donor
homology
Eμ enhancer
Cμ
Cγ1
Mouse germline Igh locus
B
317flag 9.66
317loxflag 7.93
2541flag 3.19
2541loxflag 2.70
untreated 0.037
Flag-PE
Flag-BV421
C
In vitro
% edited alleles
317flag
2541flag
B1-8hi
control
D
Chimerism
% donor CD45.2+ of B cells
HSPC edit
E
Peripheral blood CD45.2+ B cells
% IgKflag+
% edited alleles
Flow cytometry
ddPCR
F
317flag 1.24
B1-8hi 0
NANP-APC
Flag-BV421
G
anti-NANP IgG 317flag equivalents
PfCSP-ferritin
prime
boost
μg/mL
days post immunization
317flag
2541flag
B1-8hi
C57BL/6J
H
anti-NANP IgG
EC50
iGL-CIS43.D3flag
C57BL/6J

**Fig. S13. Anti-*Plasmodium falciparum* CSP antibody gene-edited HSPCs.** Related to Fig. 5 (**A**) Gene editing strategy to edit light chain flag-tagged antibodies 317, 2541 into the mouse IgH locus using 2 different linkers. (**B**) Flow cytometric analysis of flag-tagged antibody surface expression on mouse B cells *in vitro*, 2 days after editing with constructs indicated in (A). (**C**) Percentage of antibody edited alleles in mouse HSPCs at the time of transplantation by ddPCR. Each dot indicates data from an independent experiment, bars indicate median. (**D**) Chimerism among B cells 5–6 weeks after transplantation of $CD45.2^{+}$ congenic HSPCs, edited as indicated into $CD45.1^{+}$ recipients. Each dot represents a mouse. Bars indicate median ± interquartile range. Pooled data from 4 independent experiments. (**E**) Gene editing by flow cytometry and ddPCR in $CD45.2^{+}$ B cells from peripheral blood of the indicated mice 5–6 weeks after transplantation. Each dot represents a mouse. Bars indicate mean. Connecting lines indicate paired samples from the same mouse. (**F**) Flow cytometric analysis of NANP-binding and flag-tagged antibody surface expression on $CD45.2^{+}$ B cells from peripheral blood of mice 7 weeks after receiving congenic HSPCs edited with the indicated antibody. A non-flag tagged B1-$8^{hi}$ antibody was used as control. (**G**) 10 weeks after receiving $317_{flag}$, $2541_{flag}$ or control B1-$8^{hi}$ edited-HSPCs, experimental or control wildtype mice were immunized with PfCSP-ferritin nanoparticle immunogen and anti-NVDPNANP peptide serum antibody responses were measured by ELISA. Quantification of total anti-NVDPNANP IgG. Thick lines indicate geometric mean and thin lines indicate individual mice. Pool of 2 independent experiments. Concentration in $317_{flag}$ equivalents was calculated using a $317_{flag}$ recombinant antibody standard. (**H**) as in (G) but for antibody iGL-CIS43.$D3_{flag}$ immunized 6 weeks after reconstitution. Numbers in flow cytometric plots indicate percentage of cells within gate of all cells plotted

### Fig. S14

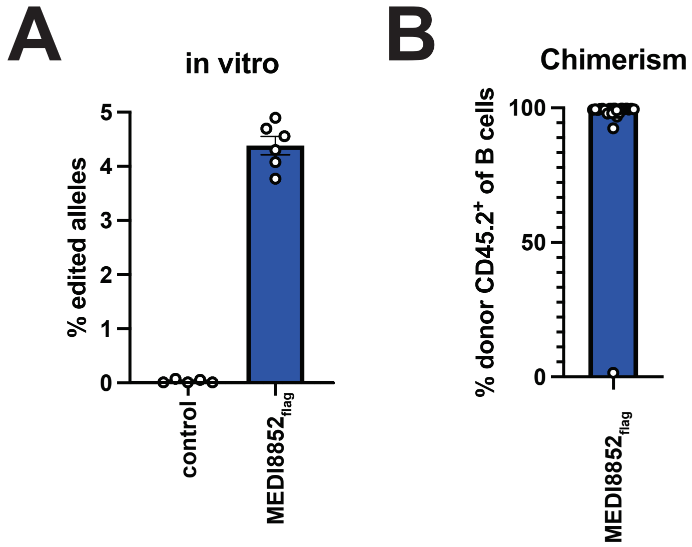
A
in vitro
% edited alleles
5
4
3
2
1
0
control
MEDI8852flag
B
Chimerism
% donor CD45.2+ of B cells
100
50
0
MEDI8852flag

**Fig. S14. Anti-Influenza bNAb gene editing.** Related to Fig. 6. (**A**) Gene-editing efficiency *in vitro* in HSPCs at the time of cell transfer by digital droplet PCR measuring target locus integration of $MEDI8852_{flag}$ using an in-out primer strategy. Controls are unedited samples. Pooled data from 3 independent experiments with 1–2 technical replicates is shown, each dot representing a technical replicate. Bars indicate mean ± SEM. (**B**) Peripheral blood chimerism among B cells 5–7 weeks after transplantation of $CD45.2^{+}$ congenic, $MEDI8852_{flag}$-edited HSPCs into $CD45.1^{+}$ recipients. Each dot represents a mouse. Bars indicate median ± interquartile range. Pooled data from 4 independent experiments.

Fig. S15

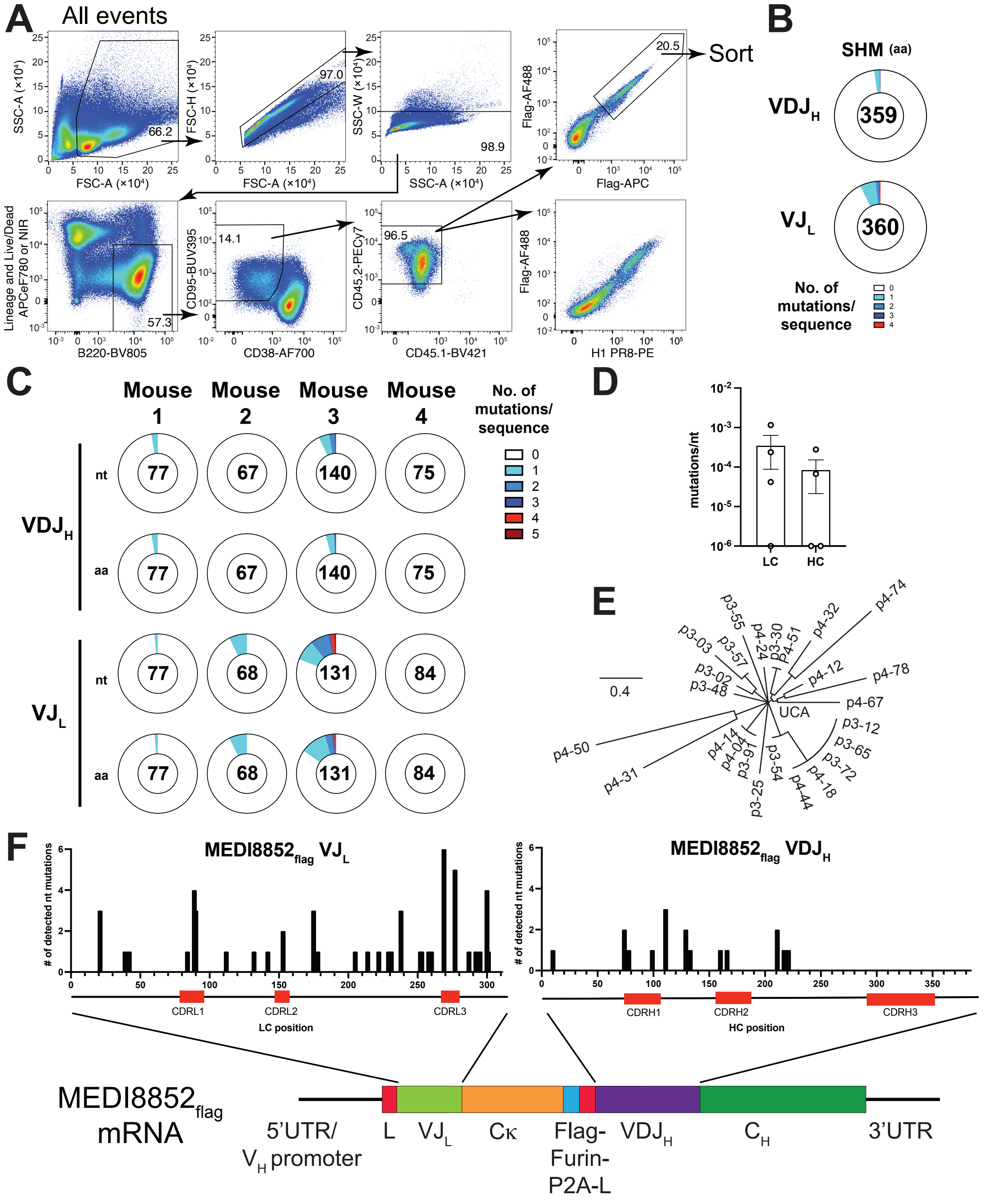

A
All events
SSC-A (×10⁴)
FSC-A (×10⁴)
66.2
FSC-H (×10⁴)
97.0
SSC-W (×10⁴)
SSC-A (×10⁴)
98.9
Flag-AF488
Flag-APC
20.5
Sort
Lineage and Live/Dead APCeF780 or NIR
B220-BV805
57.3
CD95-BUV395
CD38-AF700
14.1
CD45.2-PECy7
CD45.1-BV421
96.5
Flag-AF488
H1 PR8-PE
B
SHM (aa)
VDJH
359
VJL
360
No. of mutations/ sequence
C
Mouse 1
Mouse 2
Mouse 3
Mouse 4
No. of mutations/ sequence
VDJH
nt
77
67
140
75
aa
77
67
140
75
VJL
nt
77
68
131
84
aa
77
68
131
84
D
mutations/nt
LC
HC
E
0.4
UCA
p4-74
p4-32
p4-51
p3-30
p4-24
p3-55
p3-03
p3-57
p3-02
p3-48
p4-12
p4-78
p4-67
p3-12
p3-65
p3-72
p4-18
p4-44
p3-54
p3-25
p3-91
p4-04
p4-14
p4-50
p4-31
F
MEDI8852flag VJL
MEDI8852flag VDJH
### of detected nt mutations
CDRL1
CDRL2
CDRL3
LC position
CDRH1
CDRH2
CDRH3
HC position
MEDI8852flag mRNA
5'UTR/ VH promoter
L
VJL
Cκ
Flag-Furin-P2A-L
VDJH
CH
3'UTR

**Fig. S15. Analysis of somatic hypermutation.** Related to Fig. 6. (**A**) Flow cytometric gating strategy to sort Flag-tagged MEDI8852 expressing donor GC B cells. H1 PR8 binding of donor GC B cells is shown below. Numbers in plots indicate percentage of cells within gate of all cells plotted (**B**) Number of amino acid changes due to somatic hypermutations per sequence in heavy ($VDJ_H$, top) or light chain ($VJ_L$, bottom) regions of single-cell sorted Flag-tagged MEDI8852-expressing germinal center B cells 7 days after third immunization with H3 HK68, 322-356 days after prime. Numbers indicate total analyzed cells. Pool of 4 mice from 2 independent experiments. (**C**) as in (B) but split into individual mice and showing both nucleotide and amino acid changes. (**D**) Mutation rate in $VJ_L$ and $VDJ_H$ regions. Each dot indicates one mouse. 0 values set to $10^{-6}$ for display. (**E**) Phylogenetic tree of $VJ_L$ sequences of mouse 3. UCA, unmutated common ancestor MEDI8852 sequence. Plate and well ID are referenced (see Data S3) Scale bar indicates percentage difference. (**F**) Distribution of mutations along the $MEDI8852_{flag}$ mRNA $VJ_L$ and $VDJ_H$ regions.

### Fig. S16

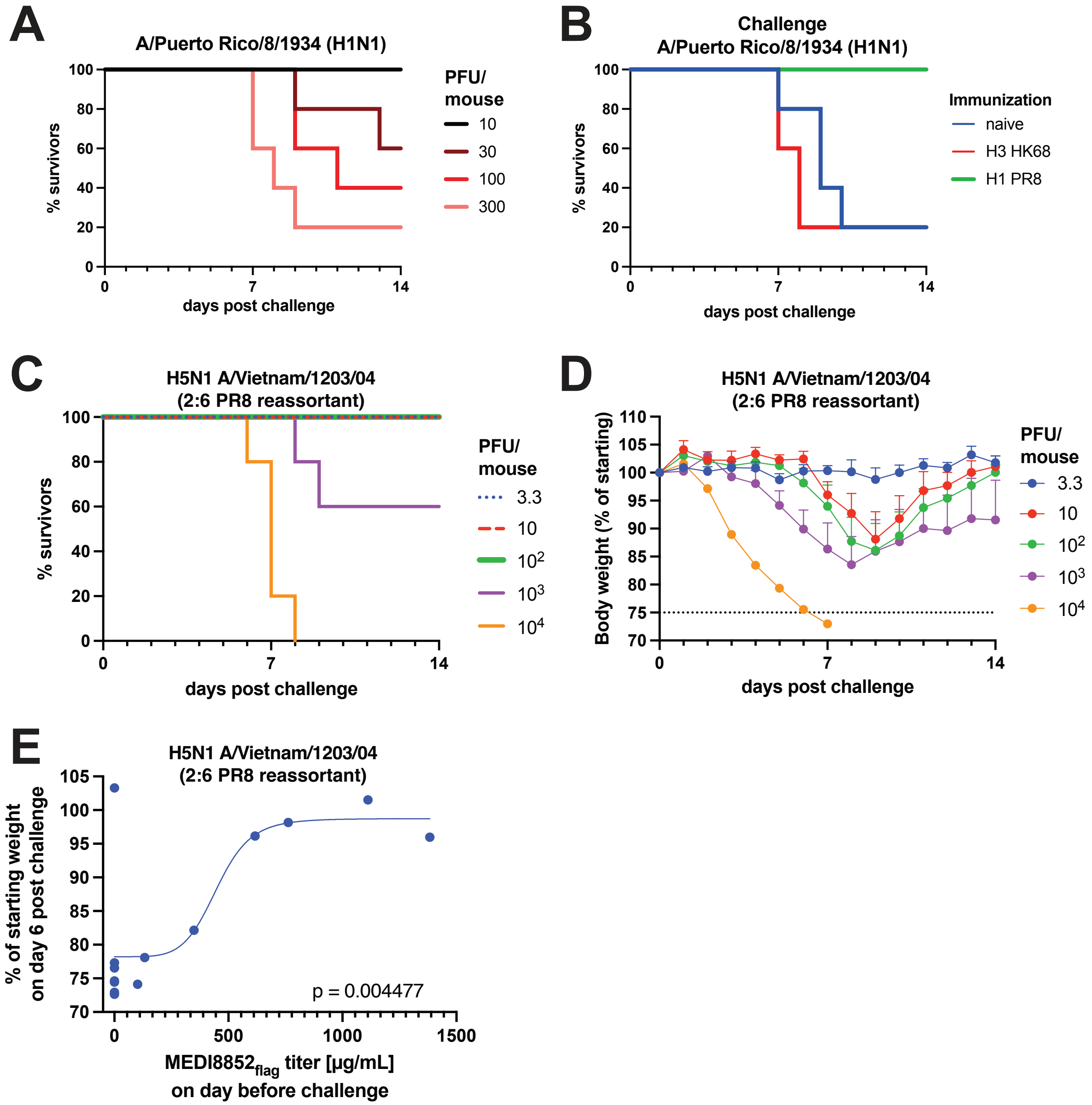
A
A/Puerto Rico/8/1934 (H1N1)
% survivors
days post challenge
PFU/
mouse
10
30
100
300
B
Challenge
A/Puerto Rico/8/1934 (H1N1)
% survivors
days post challenge
Immunization
naive
H3 HK68
H1 PR8
C
H5N1 A/Vietnam/1203/04
(2:6 PR8 reassortant)
% survivors
days post challenge
PFU/
mouse
3.3
10
$10^2$
$10^3$
$10^4$
D
H5N1 A/Vietnam/1203/04
(2:6 PR8 reassortant)
Body weight (% of starting)
days post challenge
PFU/
mouse
3.3
10
$10^2$
$10^3$
$10^4$
E
H5N1 A/Vietnam/1203/04
(2:6 PR8 reassortant)
% of starting weight
on day 6 post challenge
p = 0.004477
$MEDI8852_{flag}$ titer [μg/mL]
on day before challenge

**Fig. S16. Influenza A virus challenge models.** Related to Fig. 6. (**A**) $mLD_{50}$ determination of influenza strain PR8. Naïve C57BL/6J mice were intranasally infected with the indicated number of PFU and survival was monitored for 14 days. n = 5 per group (**B**) 2 weeks after H3 HK68 or H1 PR8 immunization or naïve control C57BL/6J mice were challenged with 5× $mLD_{50}$ of PR8. Survival was monitored for 14 days after challenge. n = 5 per group (**C**) Survival and (**D**) weight loss of naïve B6.SJL mice after intranasal challenge with the indicated dose of A/Vietnam/1203/2004 (H5N1) 2:6 PR8 reassortant with polybasic cleavage site removed to determine $mLD_{50}$. n = 5 per group (**E**) Correlation between $MEDI8852_{flag}$ titer before challenge from Fig. 6E and body weight 6 days post challenge from Fig. 6I. 5-parameter asymmetric curve fit and P value of Pearson correlation is shown. Each dot represents data from one mouse.

**Data S1. (separate file) 3BNC117 titers of Figure 1E.** Individual mouse and time point serum concentrations of 3BNC117 measured by anti-idiotypic antibody ELISA shown in Fig. 1E.

**Data S2. (separate file) Sera pooling for HIV-1 neutralization assay.** Time points for pooling and experimental details for HIV-1 neutralization assay shown in Figure S4D.

**Data S3. (separate file) Somatic hypermutations**. Individual mutations as summarized in Fig. 6 and Fig. S15.

**Data S4. (separate file) sgRNAs.** Protospacer sequences of single guide RNAs used for gene editing in this study.

**Data S5. (separate file) ddPCR primers and probes.** Nucleotide sequences of ddPCR reagents in this study.

**Data S6. (separate file) Flow cytometric reagents.** Details of flow cytometric antibodies and staining reagents used in this study.
